# *Novabrowse:* A Tool for High-Resolution Synteny Analysis, Ortholog Detection, and Gene Signal Discovery

**DOI:** 10.64898/2026.03.27.714371

**Authors:** Lennart Rikk, Ameneh Ghaffarinia, Nicholas D. Leigh

## Abstract

Accurate genome annotation remains challenging as assembly quality often exceeds annotation reliability. Resolving ambiguities of gene presence, absence, and orthology typically requires integrating two complementary lines of evidence: sequence homology between species and the conservation of gene order (i.e., synteny). BLAST remains the standard for homology detection, yet its raw output can be difficult to interpret. Existing tools address this challenge but operate at opposing scales. Alignment viewers provide detailed pairwise statistics without genomic context, while synteny tools offer chromosome-scale perspectives without sequence-level resolution. To fill this intermediate gap, we developed *Novabrowse*, an interactive BLAST results interpretation framework featuring high-resolution multi-species synteny analysis, chromosomal re-arrangement investigation, ortholog detection, and gene signal discovery. Users define a genomic region of interest in a query species and/or use custom sequences, then select one or more subject species for comparison. The pipeline retrieves query gene sequences via NCBI API integration and performs BLAST searches against each subject transcriptome or genome. Results are presented via an interactive HTML file featuring alignment statistics, chromosomal maps, coverage visualizations, ribbon plots, and distance-based clustering of high-scoring segment pairs into putative gene units. We demonstrate these capabilities by investigating *Foxp3, Aire*, and *Rbl1*, three highly conserved vertebrate genes, in the recently assembled genome of the newt *Pleurodeles waltl. Foxp3* and *Aire* have not been described in any salamander species to date, despite availability of multiple assemblies and extensive transcriptomic datasets. Using *Novabrowse*, we discovered conserved loci and gene signals for both genes in *P. waltl*, the presence of which was subsequently confirmed via Nanopore long-read RNA sequencing. In contrast, *Rbl1* analysis uncovered a chromosomal rearrangement at its expected locus with no gene signal detected, indicating a gene loss specific to *P. waltl* despite the gene’s retention in the closely related axolotl (*Ambystoma mexicanum*). Our findings demonstrate *Novabrowse*’s capacity for evidence-based evaluation of annotation artifacts, an essential capability as high-quality assemblies become more available for phylogenetically diverse species. *Novabrowse* is open source (MIT license) and freely available at: https://github.com/RegenImm-Lab/Novabrowse.

## Introduction

Advances in sequencing have dramatically improved genome assembly quality, but annotation quality has not kept pace, and resolving whether genes are unannotated or truly absent remains a fundamental challenge [1,2,3]. This has practical implications for any researcher relying on genome annotations to identify and study genes of interest, as a single missing or misannotated entry can derail an entire line of investigation. The scale of this problem is substantial. Systematic analyses have revealed that highly conserved orthologs used as benchmarks for annotation completeness (BUSCO genes) show an ∼18% false absence rate across phylogenetically diverse eukaryotes due to gene prediction failure [2]. Genes with restricted expression patterns present an additional challenge. Current annotation pipelines depend heavily on transcriptomic evidence, meaning genes expressed at low levels or restricted to specific tissues may be missed entirely when the available RNA sequencing data does not cover the relevant conditions [4]. Taken together, these challenges highlight that current methods alone cannot always resolve annotation ambiguities, and tools enabling evidence-based manual inspection of the underlying genomic context remain essential.

High-homology sequence alignments between species provide direct sequence-level evidence for resolving such ambiguities. Various computational tools have been developed for this purpose, with BLAST (Basic Local Alignment Search Tool) by Altschul et al. (1990) [5] and its variants (e.g. BLASTn, BLASTp, tBLASTn and tBLASTx) being among the most widely used. However, when dealing with complex searches involving multiple sequences or extensive alignment results, the native output of a BLAST search can become difficult to interpret. Furthermore, distinguishing true orthologs from paralogs requires additional evidence, most commonly synteny, the conservation of gene order between species [6]. However, integrating homology search with synteny analysis typically requires data processing across multiple platforms.

This fragmentation also reflects how existing tools serve complementary but distinct purposes at different levels of resolution. At one end, tools such as the NCBI BLAST web interface provide comprehensive alignment statistics and detailed sequence-level information for individual queries [7], but this granularity, focused on pairwise similarity alone, can become overwhelming when examining multiple genes simultaneously and lacks the genomic context needed to assess broader patterns of conservation. At the other end, genome-scale synteny tools such as MCScanX [8] excel at identifying large-scale collinear blocks and chromosomal rearrangements, but their broad perspective may obscure the fine-grained sequence evidence needed to evaluate individual gene annotations or resolve specific orthology questions. Researchers investigating annotation ambiguities thus face a gap between tools optimized for individual alignments and those designed for chromosome-scale block detection. This intermediate scale of analysis, where sequence-level evidence must be interpreted within a local genomic context, is currently not well served by any of the available tools.

To address these limitations, we developed *Novabrowse*, a novel BLAST results interpretation tool that combines homology search with synteny analysis while providing a middle ground between the existing approaches. For input, users specify a genomic region in a query species whose genes will be used as search sequences, and select one or more subject species whose transcriptomes or genomes will be searched for homologous matches. The tool retrieves all gene sequences from the defined query region via NCBI API [9] integration, with additional support for custom transcript or protein sequences, and performs BLASTn, tBLASTn, and/or tBLASTx searches against the subject databases using user-defined parameters. Results are filtered for isoform redundancy and presented in an interactive HTML file containing tabular alignment data alongside chromosomal localization maps for both query and all subject species. Key features include customizable multi-species comparison with interactive chromosomal localization maps, coverage visualization with identity-based color coding, high-resolution synteny analysis via interactive ribbon plot, and gene signal discovery via distance-based match clustering. *Novabrowse* delivers essential alignment information within a clear analytical framework that facilitates immediate practical interpretation of results from both genome and transcriptome database searches, while providing a foundation for downstream analysis. Here we demonstrate these capabilities through a case study involving missing gene annotations in a salamander species.

Incomplete annotations can lead to erroneous biological conclusions, such as incorrectly attributing phenotypic traits to gene absence when the relevant genes are in fact present but unannotated. Additionally, errors propagate when flawed annotations serve as templates for related species [4]. This issue becomes particularly problematic when studying species that are evolutionarily distant from established model organisms. The salamander *Pleurodeles waltl* (Iberian ribbed newt) represents one such case. As only the second chromosome-scale salamander assembly at the time of publication, it illustrates how annotation challenges intensify in phylogenetic branches with limited reference data. Despite having an available genome assembly with one of the highest contiguity achieved for a genome of this size (20.3 Gb) [10], we found that several genes expected to be present based on evolutionary conservation remain unidentified in the current annotation. This is notable because *P. waltl* has an abundance of available transcriptomic data, with over 3,200 biological samples representing diverse tissues and developmental stages publicly available on NCBI (BioSample database, accessed January 2026). However, *P. waltl* has also been demonstrated to genuinely lack certain otherwise highly conserved genes, including *Telomerase Reverse Transcriptase* (*Tert*), the catalytic subunit of telomerase [11]. Distinguishing whether these unannotated entries similarly reflect true evolutionary losses or gaps arising from the inherent limitations of current annotation pipelines to effectively process highly divergent gene sequences requires integrating multiple lines of evidence. This challenge motivated the development of *Novabrowse*. Using *Novabrowse*, we show how the integration of gene signal detection with synteny analysis enables systematic evaluation of gene presence or absence. We first demonstrate the identification of putative gene signals for *Forkhead box P3 (Foxp3)* and the *Autoimmune regulator (Aire)*, two highly conserved genes in vertebrates with critical roles in adaptive immunity that were unannotated in the *P. waltl* reference genome. The presence of these genes was subsequently verified through Nanopore-based RNA sequencing. We then apply the same methodology to *Retinoblastoma-like 1* (*Rbl1*), where *Novabrowse* revealed a chromosomal rearrangement event that preserved flanking genes but resulted in the loss of *Rbl1* itself from *P. waltl*, despite its retention in the closely related axolotl (*Ambystoma mexicanum*). Together, these case studies illustrate how *Novabrowse* facilitates evidence-based discrimination between annotation artifacts and genuine gene loss events. As genome sequencing extends to an increasing diversity of species, versatile tools that unite multiple lines of evidence for evaluating gene presence, absence, orthology, and synteny will become increasingly important for reliable genome interpretation.

## Methods

*Novabrowse* is implemented in Python 3.13.9 and can be executed in three ways: directly as a Jupyter notebook or Docker command-line, or via Docker Desktop. The pipeline utilizes NCBI BLAST+ 2.15.0 and Biopython 1.85. Installation instructions are available in the project repository at: https://github.com/RegenImm-Lab/Novabrowse

The pipeline can be divided into four sequential modules (Fig. 1). The workflow begins by defining the genomic region of interest (Target Area) in the query species and selecting subject species for comparison. BLAST search parameters are then configured independently for each subject species. The pipeline then automatically retrieves query sequences from NCBI and performs BLAST homology searches against the selected subject species databases. The results are interpreted, filtered, and assembled into an interactive HTML file.

**Figure 1.**
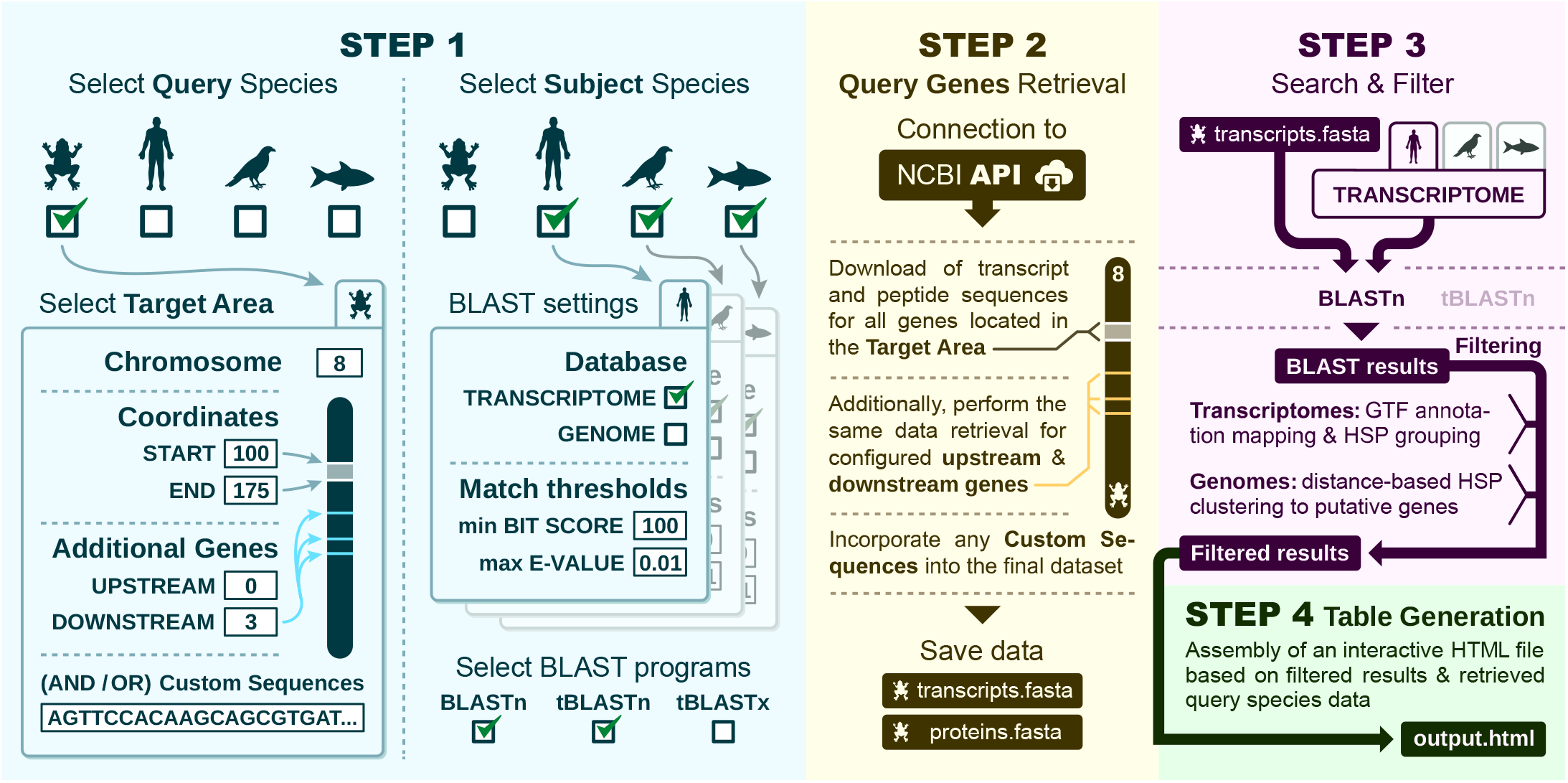
Overview of the *Novabrowse* pipeline workflow. **Step 1: Query and subject species selection with analysis parameters**. Users define the target genomic region (chromosome coordinates and optional flanking genes) in the query species, with support for custom sequences either alone or in combination with NCBI data. Subject species configuration supports database type selection (transcriptome and/or genome), BLAST algorithm choice (BLASTn, tBLASTn, and/or tBLASTx), and match filtering thresholds (minimum bit score and maximum E-value). **Step 2: Automated retrieval of query sequences via NCBI API integration**. The pipeline downloads transcript and peptide sequences for all genes located within the defined target area, along with any specified upstream and downstream flanking genes. Custom sequences, if provided, are incorporated into the dataset at this stage. Retrieved data are saved as FASTA files (transcripts.fasta and proteins.fasta) for subsequent analysis. **Step 3: BLAST search execution and results filtering**. Query sequences (transcripts and/or proteins) are searched against configured subject databases using selected BLAST algorithms (BLASTn for nucleotide-nucleotide, tBLASTn for protein-nucleotide, and/or tBLASTx for translated comparisons). For transcriptome databases, the pipeline matches BLAST hits to transcript entries in GTF annotation files to retrieve corresponding gene names, chromosomal positions, and annotation details, while consolidating multiple High-scoring Segment Pairs (HSPs) from the same transcript under the coverage column. For genome databases, chromosome identifiers are extracted from hit headers and HSPs are clustered based on a user-defined distance threshold (in base pairs) to identify putative gene units, with HSPs within the threshold distance grouped together and assigned unique identifiers (*Gene_1, Gene_2*, etc.). **Step 4: Data integration and table generation**. The pipeline parses query species FASTA files (transcripts/proteins) to extract gene metadata (names, coordinates, lengths among others) and integrates this with the filtered BLAST results from Step 3. The system subsequently maps each query gene to its ranked subject species matches and compiles the results into an interactive HTML table.

To position *Novabrowse* within the landscape of existing tools for BLAST output interpretation, we compared its feature set against five established platforms that collectively represent the major approaches to homology search, results visualization, and synteny analysis: the *NCBI BLAST* [7], *Ensembl BLAST* [12], and *UCSC BLAT* [13] web interfaces, which provide online sequence similarity searches with varying degrees of built-in results visualization; *SequenceServer* [14], an open-source web interface for running BLAST searches against user-provided databases; and *MCScanX* [8], a toolkit for synteny detection and collinearity analysis. Eight features were evaluated across all tools (Table 1).

**Table 1.**
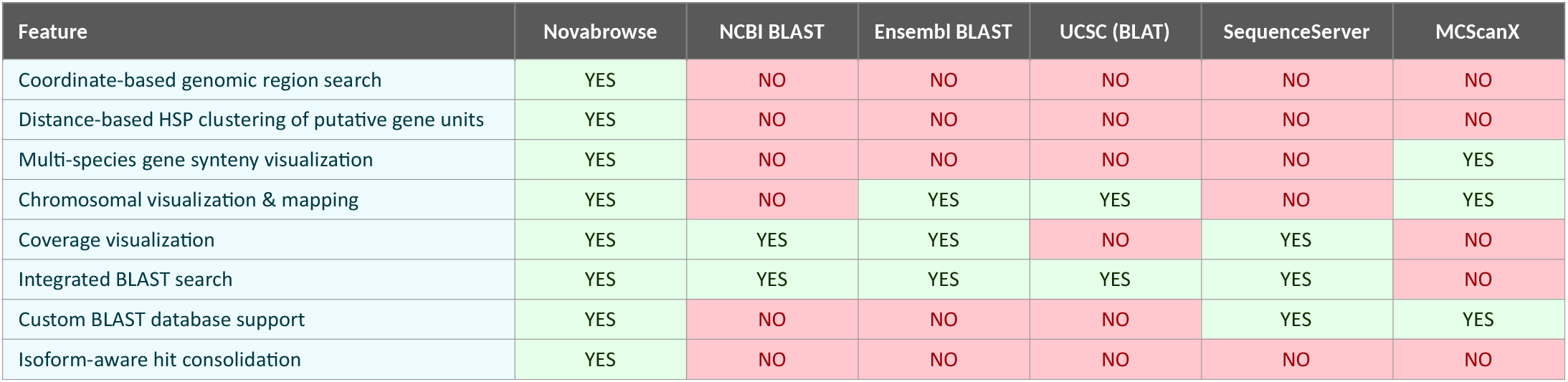
Feature comparison of Novabrowse with existing sequence analysis and comparative genomics tools. Features are evaluated as YES (fully or partially supported), NO (not supported). **Coordinate-based genomic region search** refers to the ability to define a chromosomal region by coordinates and automatically retrieve gene sequences from that region for use as search queries. **Distance-based HSP clustering of putative gene units** describes the grouping of individual high-scoring segment pairs into candidate gene units based on a user-configurable distance threshold, enabling gene signal discovery in unannotated genomes. **Multi-species gene synteny visualization** refers to the display of conserved gene order across multiple species via table and ribbon plots. **Chromosomal visualization and mapping** refers to the display of hit locations on chromosome ideograms, providing genomic context for alignment results. **Coverage visualization** describes the graphical representation of query sequence coverage by aligned HSPs. **Integrated BLAST search** indicates whether the tool natively executes BLAST or an equivalent sequence similarity search within its pipeline. **Custom BLAST database support** indicates whether users can provide their own genome or transcriptome assemblies as search targets rather than being limited to pre-built hosted databases. **Isoform-aware hit consolidation** refers to the grouping of multiple transcript isoform matches under a single gene-level entry, reducing redundancy while preserving access to individual isoform alignment statistics. Detailed evidence and references for each verdict are provided in Supplemental Table 1.

### Query Species Gene Data Retrieval

A custom Python script was developed to retrieve both protein and transcript sequences of genes from NCBI databases using the NCBI E-utilities (The Entrez Programming Utilities) API. The retrieval process requires users to first define a target genomic region by specifying the query species, chromosome name, and start/end coordinates. Each execution is limited to a single chromosome, but supports batch processing of multiple species within the same run. The script implements an adaptive search strategy that first identifies genes within the specified coordinates, then iteratively expands the search range to capture the requested number of upstream and downstream flanking genes if specified by the user. When no genes are found within the initial target region, the algorithm automatically locates the nearest upstream and downstream genes to serve as anchor points for expanding the search window.

To accommodate variations in chromosome nomenclature across different species databases, the script tests multiple chromosome format variations (e.g., “chromosome X”, “chrX”, “CHR X”) until a compatible format is identified. Users can specify protein source preferences based on NCBI accession prefixes (e.g., RefSeq curated proteins with NP_ prefixes, predicted model proteins with XP_ prefixes, among others). All transcript and protein isoforms are retrieved for each gene, provided they meet the specified protein source criteria.

The script employs error handling with an exponential backoff retry mechanism to manage network connectivity issues and NCBI server limitations. All retrieved sequences are formatted with standardized headers containing gene identifiers, chromosomal coordinates, strand information, sequence lengths, and functional descriptions. Both proteome and transcriptome datasets are generated simultaneously, with transcript-protein relationships preserved through internal mapping.

The pipeline also supports custom sequences, which can be integrated with NCBI-retrieved datasets or processed independently, allowing utilization of sequences not available in the NCBI databases.

Output from the Query Species Gene Sequence Retrieval module consists of standardized FASTA files organized hierarchically by species and chromosomal coordinates.

### Subject Species Database Construction

Subject species databases for BLAST searches are prepared from species-specific transcriptome or genome files. Any species can be incorporated as a subject database provided standardized (NCBI format) genomic annotation files (GTF (Gene Transfer Format)) and corresponding sequence files (FASTA) are available. The workflow requires manual placement of sequence files into specified directories, followed by automated processing through the BLAST+ makeblastdb utility to create indexed databases optimized for sequence searches.

### BLAST Homology Search Pipeline

A pipeline was developed to perform systematic homology searches using the retrieved query gene sequences and their isoforms against set up subject species databases. The pipeline supports three BLAST algorithms: BLASTn (nucleotide queries against nucleotide databases), tBLASTn (protein queries against nucleotide databases) and tBLASTx (translated nucleotide queries against translated nucleotide databases). Multiple BLAST search types can be executed in batch mode within a single run (though they are not run in parallel, but sequentially). The system automatically matches query sequences to appropriate BLAST algorithms (e.g., protein sequences are used for tBLASTn searches while transcript sequences are used for BLASTn and tBLASTx searches).

The pipeline also enables searches against multiple subject species using the same query information within a single execution (again, the searches are not run in parallel, but sequentially). Users can configure which species to include for each execution and customize search parameters on a per-species basis, including E-value thresholds and bit score cutoffs. This is useful because while bit scores are database size independent, E-values are relative to database size, enabling users to apply appropriate filtering thresholds that account for the varying sizes of different species’ databases, ensuring consistent and meaningful comparisons across subject species with genomes or transcriptomes of different sizes.

### BLAST Results Processing

The BLAST search output files undergo automated parsing to extract homologous sequences and map them to annotated genes. The parsing algorithm processes standard BLAST text format files by first automatically detecting the BLAST type, then identifying query sequences with hits and extracting match identifiers, scores, E-values and alignment coordinates from each High-scoring Segment Pair (HSP).

For transcriptome-based searches, gene annotation is performed using species-specific GTF (NCBI format) files containing chromosomal coordinates, gene identifiers, and functional descriptions. The system matches BLAST hit identifiers to transcript entries in the GTF files, retrieving corresponding gene names, chromosomal positions, and annotation details. Multiple HSPs from the same transcript mapping to the same query gene are consolidated. This means that if a query hits the same transcript multiple times at different positions, all those HSPs are collected under that single transcript ID and output as one in the results with all HSPs listed under the coverage column.

For genome-based searches, GTF annotation files are not used, as the objective is to identify unannotated gene signals. Instead, the system first extracts chromosome identifiers directly from BLAST hit headers by recognizing standardized accession patterns (NC_, NW_, Chr*, etc.), then implements distance-based clustering to group nearby HSPs into novel putative gene units based on their proximity in the genome [15][16]. The algorithm processes all HSPs for each query-subject pair by first sorting them according to their subject sequence positions, then iteratively grouping HSPs whose boundaries fall within a user-defined distance threshold (specified in base pairs). This clustering works by calculating the minimum distance between any position in a candidate HSP and all positions in existing groups, if this minimum distance is less than or equal to the threshold, the HSP is added to that group (Fig. 2), otherwise, it initiates a new group.

**Figure 2.**
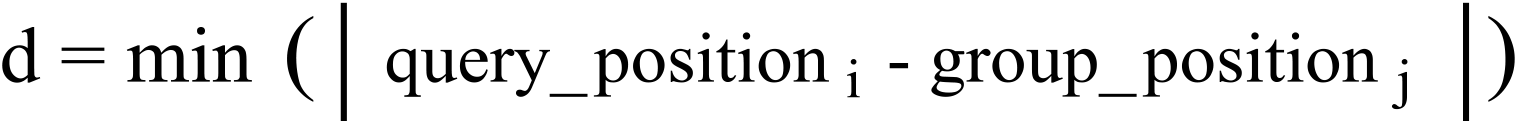
Distance calculation formula where query_positioni represents any coordinate (start or end) from the candidate HSP and group_positionj represents any coordinate from HSPs already in the group. If d ≤ user-defined distance threshold, the candidate HSP is added to the group which forms a putative gene unit.

Each resulting HSP cluster is treated as a potential gene unit and assigned a unique identifier (*Gene_1, Gene_2*, etc.), with the cluster’s boundaries defined by the minimum and maximum subject positions across all constituent HSPs. Each identified cluster, representing a putative gene unit, is documented in the coverage field as an array containing all its constituent HSPs, with each HSP preserving its individual alignment coordinates, scores, E-values, and identity percentages. This approach enables discovery of gene signals by identifying regions of concentrated homology that could represent functional gene units, with the distance threshold parameter allowing users to adjust clustering sensitivity based on the expected intergenic distances in their target species.

Both processing paths, transcriptome-based (annotation-dependent) and genome-based (annotation-independent), converge on a common tab-delimited output file but contribute distinct types of information. Transcriptome results include matched gene IDs and names retrieved from GTF annotations alongside transcript IDs, whereas genome results contain only the *Novabrowse* assigned HSP cluster identifiers (*Gene_1, Gene_2*, etc.) and the genomic coordinates defining each putative gene unit, with no cross-referencing to existing annotations. Both paths record chromosome identifiers, alignment scores, E-values, identity percentages, subject and query lengths, full query descriptors, and coverage arrays detailing all constituent HSPs. This modular approach allows users to selectively process subsets of their data while maintaining a complete audit trail of all processing attempts, ensuring both flexibility in analysis design and transparency in result generation.

### Results Visualization

The filtered BLAST results are processed to generate interactive HTML files for comparative analysis. The visualization pipeline integrates query species sequence data with BLAST matches from one or more subject species to create a table of the results along with chromosomal visualizations of relevant gene positions. All interactive features are implemented using JavaScript, enabling dynamic filtering, data manipulation, and visualization updates directly in the web browser without requiring server-side processing.

The table generation process begins by reading query species transcriptome FASTA files to obtain gene identifiers, transcript IDs, chromosomal positions (start/end coordinates), DNA and mRNA sequence lengths, chromosome assignments, and gene names. Simultaneously, the pipeline processes subject species BLAST results, extracting matched gene identifiers, alignment coordinates on subject chromosomes, sequence lengths, BLAST scores, E-values, identity percentages (both total and local), coverage metrics, best local similarity scores, query lengths, and HSP details. User-defined filtering parameters, including minimum score thresholds and maximum E-value cutoffs, are applied during data integration for each subject species.

The HTML table is constructed with query genes displayed as rows in the first column and corresponding genes matched from subject species in adjacent columns (Fig. 3). When multiple isoforms of a query gene exist, the isoform producing the highest-scoring alignment is selected for inclusion in the table. When multiple subject species genes match a single query gene, the system ranks them by alignment score for transcriptome databases, or by coverage percentage for genome databases, designating the highest-ranking match as primary and all others as secondary. Users can configure the table to include either all matches (both primary and secondary) for comprehensive homology analysis, or only primary matches for a more streamlined view with improved rendering performance.

**Figure 3.**
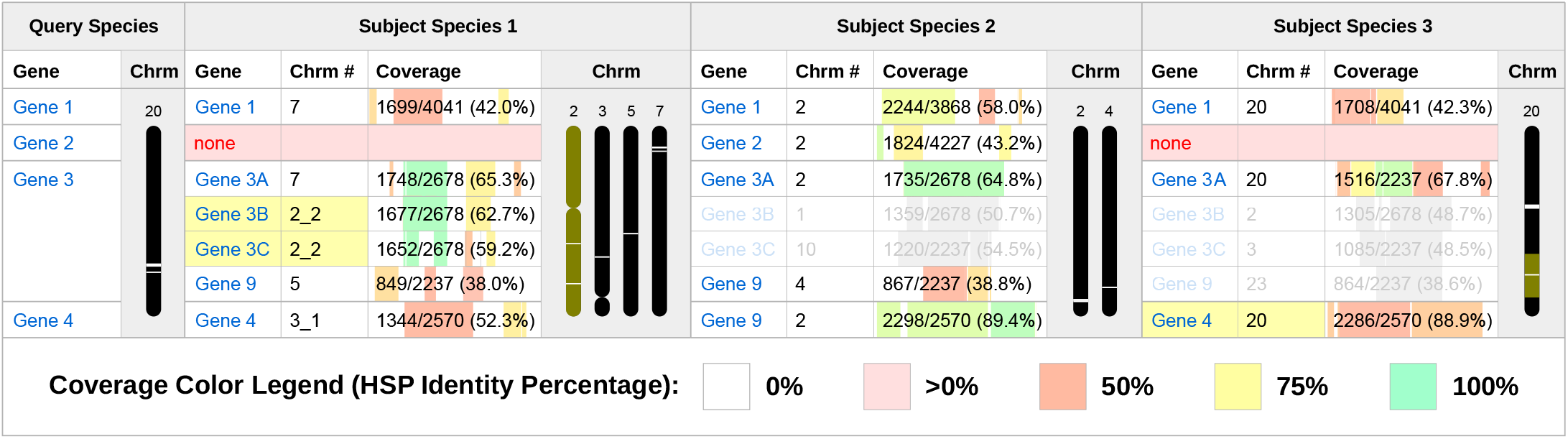
*Novabrowse* table layout example. The Query Species column (left) lists genes used as queries for BLAST searches. The “Coverage” column shows alignment coverage as colored bars positioned along the query sequence, indicating where HSPs align, with values shown as both percentages and absolute lengths (total unique subject HSP regions length/query transcript length). Coverage bars are color-coded by HSP identity percentage according to the legend shown below. The “Chrm” column displays visualized chromosomes showing relative gene positions, with chromosome heights normalized across the table. The “Chrm #” column indicates chromosomal locations, with underscores denoting different chromosome arms (e.g., 2_2, 3_1). Yellow backgrounds indicate genes highlighted through the coordinate-based area highlighting feature: selected chromosomal regions are highlighted in yellow on the chromosome visualization, and all genes mapping to those regions are also marked with yellow backgrounds in the gene table. Red “none” entries indicate genes without matches in that species, while grayed-out genes indicate matches found but excluded by the chromosomal filtering feature. Note: Gene names in the figures are displayed as they appear in NCBI databases, while species-appropriate nomenclature conventions are applied throughout the text.

Each gene match in the subject species columns shows the number of transcript isoforms found, with only the top-scoring isoform info shown by default. Clicking the transcript count button reveals all matched isoforms with their individual alignment statistics (coverage, identity percentages, scores, and E-values). When the same isoform has multiple matches per gene (different segments), the system shows only the best-scoring segment information. However, the coverage visualization ensures no alignment information is lost by displaying all aligned segments per isoform as bars positioned along the query sequence, showing exactly how each section aligns to the query gene.

Coverage is calculated as the proportion of the query sequence that aligns to the subject sequence across all HSPs, using the formula shown in Fig. 4. The calculation accounts for overlapping HSP regions by storing each aligned position in a set, merging any overlapping intervals before computing coverage, ensuring accurate representation without double-counting overlapped segments. These coverage values are displayed as positioned bars along the query sequence, visually indicating where HSPs align (Fig. 3). The coverage bars are color-coded by HSP identity percentage to convey alignment quality. This approach allows users to simultaneously assess the extent of sequence similarity (through coverage percentage), the distribution of aligned regions (through the positioned bars), and quality of each region (color-coding). Such visualization can be valuable for identifying partial homologs, domain-specific matches, or genes with conserved regions interspersed with divergent sequences.

**Figure 4.**
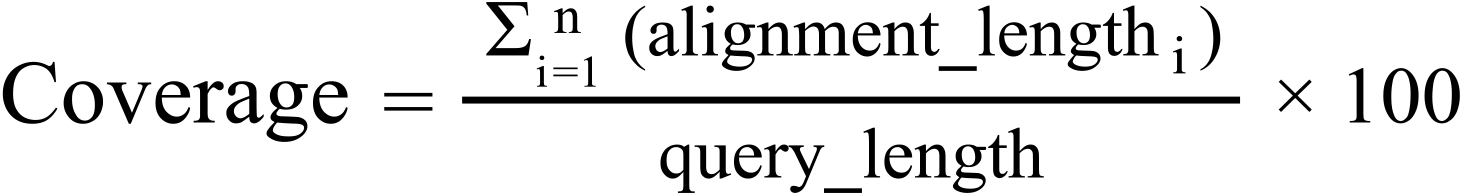
**Coverage calculation formula** where alignment_length_i_ is the length of the i-th HSP segment. The result is expressed as a percentage.

Chromosomal localization is implemented through interactive chromosome maps positioned within species-specific columns. The coordinate system accommodates both single chromosomes and multi-arm configurations (p and q arms), with automatic coordinate transformation to maintain proper relative positioning across chromosome arms. This dual-format support is necessary because genome assemblies may represent chromosomes either as complete units or as separate arms depending on assembly rather than on actual biological structure (i.e., a single-unit representation does not necessarily mean only one arm is shown, rather, it typically means that both arms were combined into a single entry). Gene positions are calculated relative to total chromosome length and displayed as positioned elements along chromosome representations (Fig. 3). Chromosome visual height can be normalized to identical scales across species or displayed at relative scales based on actual chromosome lengths.

Advanced filtering capabilities include multiple operational modes: gene name-based filtering allows users to specify subsets of genes for analysis: “At least 1 match” mode displays only genes with at least one match in any of the subject species; “Full conservation” mode shows only genes with matches across all of the subject species: “Hide 2nd match genes” mode simplifies display by showing only the highest-scoring (primary) match per gene.

The interface allows toggling visibility of both query and individual subject species columns, as well as drag-and-drop reordering of subject species to facilitate comparative analysis. Individual data columns can also be toggled independently, including gene identifiers (query and subject), genomic coordinates (start, end, chromosome number), length metrics (DNA length percentage, mRNA length percentage, query length), alignment quality parameters (best total identity percentage, best local identity percentage, coverage, number of HSPs), statistical measures (score, E-value) and chromosome columns, allowing users to customize data visualization according to analytical requirements.

Coordinate-based area filtering facilitates focused analysis of specific genomic regions through highlighting and retention tools. The “Highlight” feature emphasizes genes within user-defined coordinate ranges by applying visual markers in both the table and visualized chromosomes, while the “Keep” filter provides more restrictive control by displaying only genes that fall within specified coordinate ranges on selected chromosomes (Fig. 3). When Keep is activated, genes outside the defined spans are hidden from view, and entire chromosomes not selected through the checkbox system are removed from the display, isolating the genomic regions of interest for detailed examination.

As filters are applied, the system provides continuous statistical tracking with real-time updates of match counts, species-specific match tallies, and highlighted gene counts.

The ribbon plot algorithm determines the relative positions of genes across all displayed species and creates smooth, curved pathways linking putatively homologous genes on the chromosomes. Users can achieve high-resolution synteny visualization by hovering over individual genes, which highlights the corresponding ribbon across all species. Additionally, visual clutter can be reduced by interface functionality that allows users to customize the style of each ribbon and selectively display ribbon connections for specific genes only.

Export capabilities generate SVG files containing the complete table structure with all data columns, chromosome maps, and active ribbon plot visualizations. These files are ready for publication or further editing in vector graphics programs (e.g., Inkscape, Adobe Illustrator, and others).

### Data & BLAST Parameters

Query sequences were retrieved from NCBI databases. Human *FOXP3* transcript sequences were obtained from chromosome X (coordinates 49,250,521–49,264,710; assembly GCF_000001405.40). For *Xenopus tropicalis foxp3*, sequences were retrieved from chromosome 8 (coordinates 25,943,647–25,959,613; assembly GCF_000004195.4). Human *AIRE* sequences were obtained from chromosome 21 (coordinates 44,285,876–44,298,648; assembly GCF_000001405.40). *X. tropicalis aire* sequences were obtained from chromosome 8 (coordinates 126,425,535–126,448,861; assembly GCF_000004195.4). *A. mexicanum Rbl1* sequences were obtained from chromosome 3p (coordinates 889,108,772–890,221,502; assembly GCF_040938575.1). Human *RBL1* sequences were obtained from chromosome 20 (coordinates 36,996,349 –37,095,997; assembly GCF_000001405.40). For high-resolution synteny analysis of the expected *P. waltl Rbl1* locus, 60 genes were retrieved from *P. waltl* chromosome 7 (All genes falling to 411,635,276– 447,535,149 + 21 genes upstream and 7 genes downstream of those coordinates); assembly GCF_031143425.1). All retrieved sequences, including upstream and downstream flanking genes, were filtered to retain only protein-coding genes with NCBI RefSeq curated protein annotations (NP_ prefix) or predicted protein model annotations (XP_ prefix).

BLAST homology searches were performed using tBLASTx algorithm against subject species databases downloaded from NCBI, including *Homo sapiens* (assembly GCF_000001405.40), *Pleurodeles waltl* (assembly GCF_031143425.1), *Mus musculus* (assembly GCF_000001635.27), *X. tropicalis* (assembly GCF_000004195.4), *Lepisosteus oculatus* (assembly GCF_040954835.1) and *A. mexicanum* (GCF_040938575.1), *Anolis carolinensis* (assembly GCF_035594765.1) and *Gallus gallus* (assembly GCF_016699485.2). An E-value threshold of 1×10^−10^ and a minimum bit score of 0 were applied for all searches. For genome-based searches targeting unannotated regions, a distance-based clustering threshold of 1,200,000 bp was used to group HSPs into putative gene units.

### Nanopore Sequencing and transcriptome generation

To obtain full-length Foxp3 and Aire transcripts, long-read Nanopore cDNA sequencing was performed on thymus tissue collected from nine pre-metamorphic (stage 43-45) wild type P. waltl individuals. Immediately after dissection, each thymus was snap-frozen in liquid nitrogen. Thymus tissues were pooled to generate a single sample and pulverized using a disposable pestle while keeping the sample tube on dry ice. TRI Reagent (Sigma, T9424) was added, and homogenates were stored at −80 °C until RNA extraction. Total RNA was then extracted according to the manufacturer’s instructions. Spectrophotometric analysis using the NanoPhotometer NP80 (Implen GmbH, Germany) indicated an RNA concentration of 138.92 ng/µL, with A260/A280 and A260/A230 absorbance ratios of 1.89 and 1.68, respectively. Fluorometric quantification using the Qubit RNA HS Assay Kit (Invitrogen, Q32852) measured an RNA concentration of 166 ng/µL.

Full-length cDNA libraries were prepared from 500 ng total RNA using the cDNA-PCR Sequencing Kit V14 (SQK-PCS114; Oxford Nanopore Technologies) according to the manufacturer’s instructions. Reverse transcription and strand-switching were used to generate full-length cDNA. cDNA was PCR-amplified to enrich full-length transcripts using rapid attachment primers (15 cycles; 8 min extension). The amplified cDNA library was quantified using the Qubit dsDNA HS Assay Kit (Invitrogen, Q32851), and the concentration of the library was 199 ng/µL. Library quantification and quality assessment were performed using an Agilent 2100 Bioanalyzer with a High Sensitivity DNA analysis kit (Agilent). cDNA libraries were run in duplicate, exhibiting a fragment size distribution of approximately 200–7252 bp, with an average size of 1763 bp. The average library concentration across duplicates was 1.614 ng/µL. A total of 50 fmol of the library was ligated with adapters and loaded onto a primed flow cell (FLO-MIN114; Oxford Nanopore Technologies). Sequencing was performed on a MinION Mk1B device (Oxford Nanopore Technologies) for 72 h. Reads were basecalled using model dna_r10.4.1_e8.2_400bps_sup@v4.3.0.

Output FASTQ files were analyzed with the Oxford Nanopore Technologies wf-transcriptomes Nextflow workflow v1.6.1 using the *Pleurodeles waltl* NCBI reference assembly (GCF_031143425.1). The corresponding reference genome FASTA and gene annotation GTF were downloaded from NCBI, decompressed, and provided to the workflow via --ref_genome and --ref_annotation. Reads were mapped to the reference genome within wf-transcriptomes using Minimap2 in splice-aware mode, and alignments were sorted and indexed with Samtools. Because the genome contains reference sequences exceeding the BAM.bai indexing range, the wf-transcriptomes repository was forked and modified to generate CSI indices by replacing.bai with.csi in main.nf and reference_assembly.nf (https://github.com/RegenImm-Lab/wf-transcriptomes-forked). The workflow was run in cDNA mode (--cdna_kit SQK-PCS114) on the basecalled FASTQ directory under the Singularity profile with 48 threads and --igv enabled, producing genome-aligned transcript models and a merged transcriptome that included transcripts assigned to unannotated genomic regions (code=u).

### Independent identification and genomic localization of *Foxp3* and *Aire* transcripts

Transcripts labeled code=u (unannotated) were extracted from the wf-transcriptomes merged transcriptome and translated in silico by predicting open reading frames with TransDecoder.LongOrfs (TransDecoder v5.7.1 [17]). The longest ORF per transcript was retained as a peptide sequence. The peptide set was screened by BLASTp against a locally downloaded UniProtKB/Swiss-Prot database using -max_target_seqs 1 and tabular output (-outfmt 6) to identify candidate transcripts with best matches to *FOXP3-* and *AIRE*-related proteins. This screen identified a candidate *Foxp3* translated transcript (***fastq_pass_batch_114*.*23*.*1***) with best match to human *FOXP3* and a candidate *Aire* translated transcript (**fastq_pass_batch_129.277.1**) with best match to human *AIRE*. Candidate transcript nucleotide sequences were then aligned to the *Pleurodeles waltl* genome assembly (GCF_031143425.1) using megablast.

### *Novabrowse* based genomic localization and identification of *Foxp3* and *Aire* transcripts

Nanopore-derived transcripts were analyzed using *Novabrowse* to establish the identity and genomic localization of *Foxp3* and *Aire* in *P. waltl*. Transcripts identified from the Nanopore dataset were incorporated into *Novabrowse* as custom query sequences (*FOXP3_nanopore* and *AIRE_nanopore*) and analyzed using tBLASTx searches against the *P. waltl* genome assembly (GCF_031143425.1). *Novabrowse*’s distance-based HSP clustering algorithm was used to group spatially proximal high-scoring segment pairs into putative gene signals employing a clustering threshold of 1,200,000 bp.

For synteny analysis validating the identified gene signals, upstream and downstream flanking genes around the *FOXP3_nanopore* and *AIRE_nanopore* loci were obtained from the same assembly (GCF_031143425.1).

## Results

### Absent Ortholog Detection

Upon manual inspection of the *P. waltl* RefSeq annotation, we noted the lack of annotations for critical immune genes *Foxp3* and *Aire*. To confirm that *Foxp3* and *Aire* are not misannotated in *P. waltl*, we utilized *Novabrowse*’s integrated pipeline to perform tBLASTx searches against the *P. waltl* transcriptome using human *FOXP3* and *AIRE* transcripts as query sequences. We then employed *Novabrowse*’s synteny analysis features to attempt to distinguish true orthologs from paralogs.

The search for *Foxp3* identified 38 hits spanning chromosomes 1-12 (excluding chromosome 8) with coverage ranging from 7.8% to 22.4% (Fig. 5A). Most high-confidence matches corresponded to *Forkhead box* gene family members, with some having established gene symbols and others bearing systematic Loc identifiers (such as *Loc138259942*, putatively a *Forkhead box protein A4-A-like* gene on NCBI, among others), all containing conserved Forkhead Box domains. However, matches with coverage outside the conserved Forkhead domain alone corresponded exclusively to genes with established symbols, while putatively annotated Loc genes showed only single-domain matches with lower coverage (7.8-9.4%). This pattern suggests that while numerous Forkhead family members exist throughout the *P. waltl* genome, none represent obvious *Foxp3* orthologs based on the sequence similarity alone, necessitating synteny analysis to identify an ortholog.

**Figure 5.**
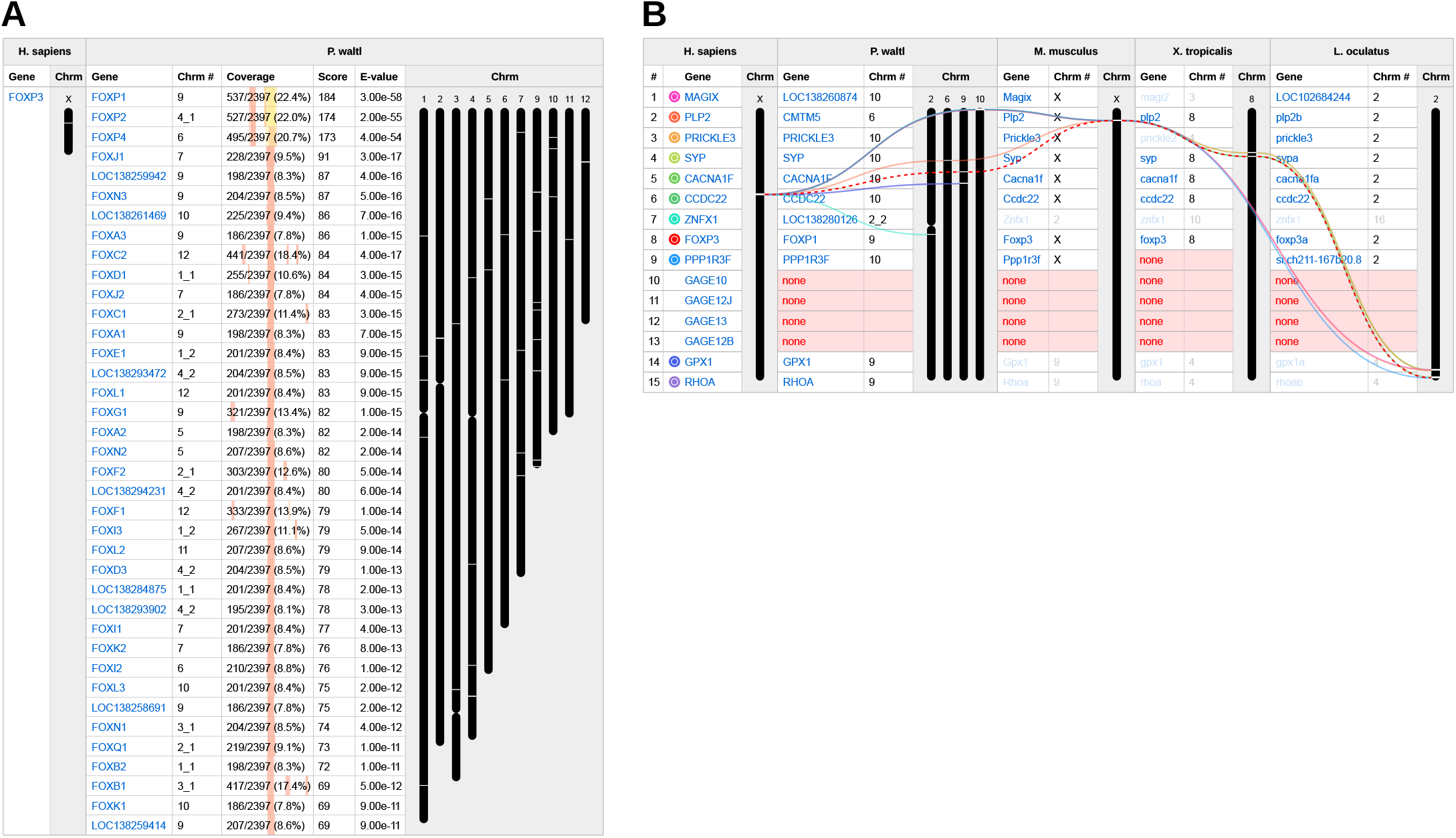
*Foxp3* ortholog detection and synteny analysis in *P. waltl* using *Novabrowse*. **(A)** tBLASTx search results of human *FOXP3* against the *P. waltl* transcriptome, showing homologous matches. Chromosome sizes are scaled proportionally to their actual lengths. **(B)** Syntenic relationships of human *FOXP3* and 14 flanking genes across human (*H. sapiens*), Iberian ribbed newt (*P. waltl*), mouse (*M. musculus*), African clawed frog (*X. tropicalis*), and spotted gar (*L. oculatus*). The analysis was performed using tBLASTx searches with human gene sequences as queries against each target species’ transcriptome, followed by chromosomal filtering to display only chromosomes with *Foxp3* matches (excluding *P. waltl* where all chromosomes with matches are shown). Query genes are marked with colored circles corresponding to ribbon colors for tracking conservation patterns. The red dashed ribbon indicates *FOXP3*. Chromosome heights are normalized to equal heights. See Fig. 3 for further layout interpretation.

For *Foxp3* synteny analysis, we examined seven upstream and seven downstream genes flanking human *FOXP3*. We then compared the positions of these genes across *P. waltl* and three additional vertebrate species where *FOXP3* has been annotated: mouse (*Mus musculus*), frog (*Xenopus tropicalis*), and spotted gar (*Lepisosteus oculatus*). The *Novabrowse* ribbon plot feature combined with its chromosomal filtering functionality, which allows selective display of matched genes within user-defined chromosomes, revealed strong conservation of the *Foxp3* genomic neighborhood across the selected taxa (Fig. 5B). The results show that most flanking genes have maintained their relative chromosomal positions, a pattern consistent with evolutionary preservation of functionally similar gene clusters [18]. Notably, in *P. waltl*, the majority of genes syntenic to the human *FOXP3* locus were found clustered on the top of chromosome 10, indicating that this region represents the orthologous genomic neighborhood where *Foxp3* would be expected to reside based on synteny conservation. However, despite the preservation of this syntenic block, no *FOXP3* orthologs or paralogs were identified within this chromosomal region in *P. waltl* (Fig. 5A), suggesting a possible annotation gap that warranted further investigation.

Having established the syntenic context for *Foxp3*, we next applied the same approach to investigate the second unannotated immune gene, *Aire*. BLAST analysis of human *AIRE* transcript against the *P. waltl* transcriptome identified nine potential matches with coverage ranging from 5.1% to 14.8% (Fig. 6A). The highest-scoring match was *Loc138265795* on chromosome 11 (annotated as “nuclear body protein SP140-like” on NCBI), followed by several immune-related genes such as *Chd4, Trim24*, and *Chd5* among others, that share functional domains with *AIRE*. To validate *Loc138265795* as a potential *AIRE* ortholog, we performed synteny analysis using the same approach as for *FOXP3*. Examination of human *AIRE* and seven upstream and seven downstream genes revealed no conserved syntenic relationships around *Loc138265795*, with identified matches to the flanking genes scattered across different chromosomes in *P. waltl* (Fig. 6B). This lack of syntenic support suggests that *Loc138265795* is unlikely to be the true *AIRE* ortholog. Given this, we examined how *AIRE* synteny conservation varies across vertebrate species representing different evolutionary distances: mouse (*Mus musculus*), lizard (*Anolis carolinensis*), chicken (*Gallus gallus*), African clawed frog (*Xenopus tropicalis*), and spotted gar (*Lepisosteus oculatus*) (Fig. 6B). The expanded analysis using *Novabrowse’s* gene highlight feature, which allows selective highlighting of matched genes within user-defined genomic coordinates, revealed that synteny conservation around *AIRE* orthologs becomes progressively weaker as evolutionary distance from humans increases. This suggests that phylogenetically closer species might provide a more optimal syntenic reference for *P. waltl*.

**Figure 6.**
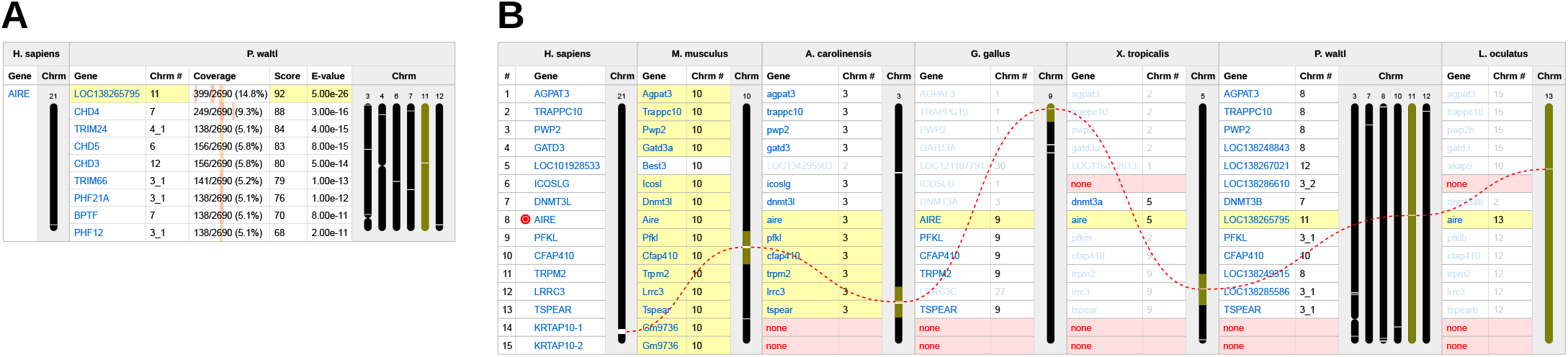
*Aire* ortholog search and cross-species synteny comparison in *P. waltl* using *Novabrowse*. **(A)** tBLASTx search results of human *AIRE* against the *P. waltl* transcriptome, showing homologous matches. Chromosome heights are normalized to equal heights. **(B)** Syntenic relationships of human *AIRE* and 14 flanking genes compared across human (*H. sapiens*), mouse (*M. musculus*), lizard (*A. carolinensis*), chicken (*G. gallus*), African clawed frog (*X. tropicalis*), Iberian ribbed newt (*P. waltl*), and spotted gar (*L. oculatus*). The analysis was performed using tBLASTx searches with human gene sequences as queries against each target species’ transcriptome, followed by chromosomal filtering to display only chromosomes with a match to *AIRE* (filtering not applied to *P. waltl*). Query genes are listed in the leftmost column. The red dashed ribbon indicates *AIRE*. Chromosome heights are normalized to equal heights. See Fig. 3 for further layout interpretation.

Given that *X. tropicalis* is the closest amphibian species with annotated *aire* in the NCBI dataset, we performed an additional analysis using its *aire* gene as the query. *X. tropicalis aire* ortholog search yielded similar results to the human *AIRE* query, with *Loc138265795* again producing the highest ranking alignment (Supplemental Fig. 1). Synteny analysis of 60 genes flanking *X. tropicalis aire* (30 upstream and 30 downstream) revealed that 45 mapped to *P. waltl* chromosome 11 within a syntenic block that shows significant expansion relative to *X. tropicalis* (Fig. 7). Notably, *Loc138265795* was the highest-scoring match for both *X. tropicalis aire* and its immediate upstream neighbor *sp110*.

**Figure 7.**
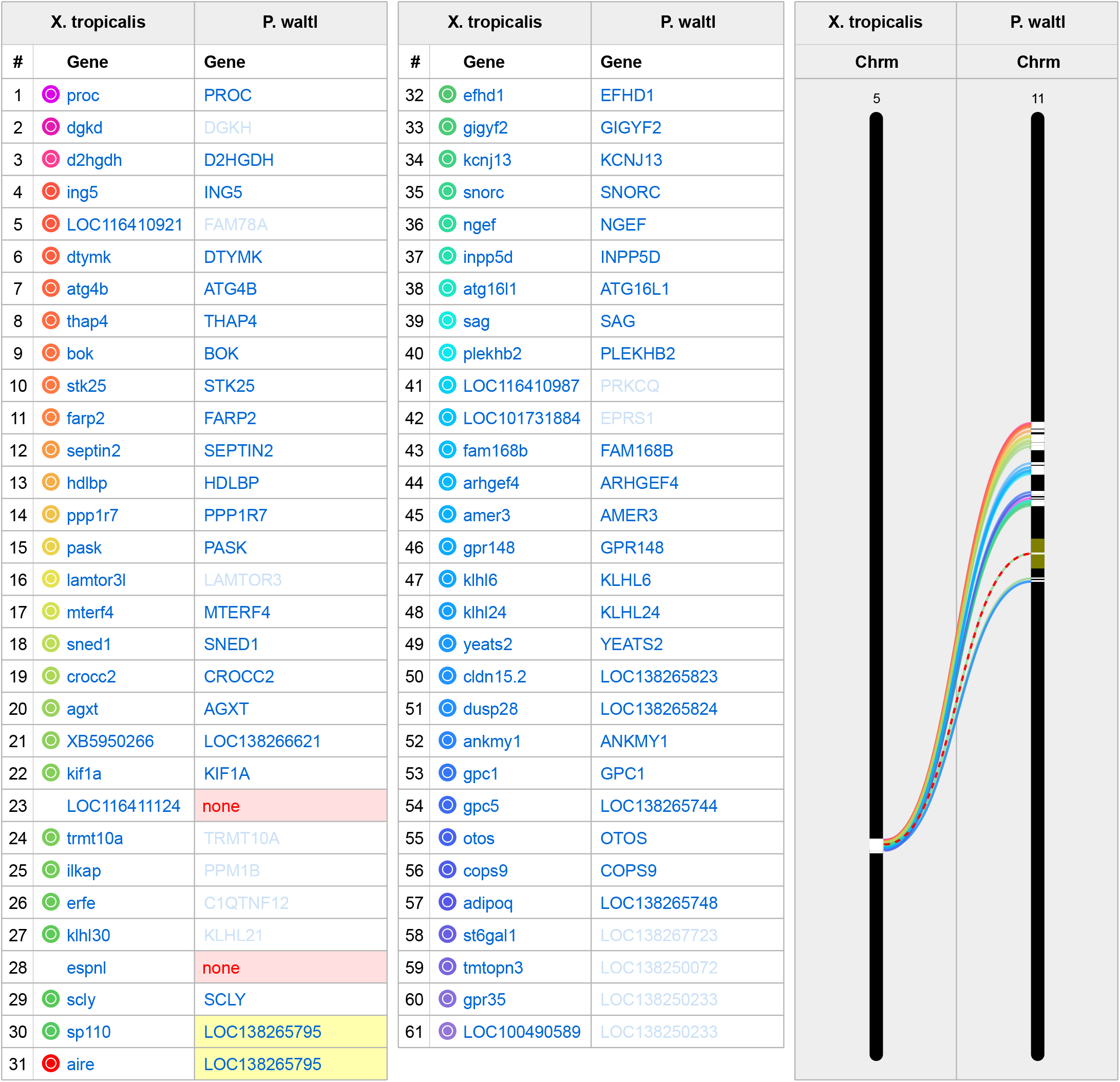
Synteny analysis of the *aire* genomic region between *X. tropicalis* and *P. waltl.* *Novabrowse* visualization examining syntenic relationships between 60 genes flanking *X. tropicalis aire* (30 upstream, 30 downstream) and their matching homologs in *P. waltl*. The analysis was performed using tBLASTx searches with *X. tropicalis* gene sequences as queries against the *P. waltl* transcriptome, with subsequent chromosomal filtering applied to display only *P. waltl* chromosome 11, where the majority of syntenic matches clustered. Due to space constraints, the gene list is split into two sections (genes 1-31 and 32-60), with each section showing *X. tropicalis* query genes and their *P. waltl* matches. Query genes are numbered sequentially and marked with colored circles corresponding to ribbon colors. Curved ribbons connect orthologous genes between species. The *aire* gene (#31, red circle) and its corresponding match in *P. waltl* are connected with a red dashed ribbon. Chromosome heights are normalized to equal heights. See Fig. 3 for further layout interpretation.

Downstream analysis of the top *P. waltl Aire* candidate (*Loc138265795*) revealed that it has five protein isoforms annotated on NCBI, three named “nuclear body protein SP140-like protein” and two named “sp110 nuclear body protein-like” (Supplemental Table 2). Based on this, *Loc138265795* is most likely an ortholog of *X. tropicalis sp110* rather than *aire*, with the alignment to *aire* likely reflecting shared functional domains.

This discrepancy between synteny conservation and gene absence for both *Foxp3* and *Aire* suggests two possible scenarios: that these genes have been lost from *P. waltl* during evolution while the flanking genes have remained, or alternatively, that these genes are present but have remained unannotated, presumably due to lack of transcriptomic evidence, in current genome assemblies. In support of the latter, searches against available long-read transcriptomic datasets from spleen, brain, and blastema tissues also failed to identify matches for either gene (data not shown).

### Gene Signal Identification

The synteny analyses described above revealed that genomic regions orthologous to *FOXP3* and *AIRE* loci are conserved in *P. waltl*, yet no transcripts corresponding to these genes were identified in the transcriptome database. We reasoned that, since gene annotations are largely derived from RNA-seq data mapped to the genome, genes expressed at low levels or restricted to specific tissues may not be captured if those conditions are not sampled. As a result, such genes would be absent from the current datasets. To investigate whether these genes might nonetheless be present in the genome, we employed *Novabrowse* to perform tBLASTx searches directly against the *P. waltl* genome assembly.

For genome-based searches, *Novabrowse* implements distance-based clustering to group High-scoring Segment Pairs (HSPs) into putative gene units, as genomic searches lack the gene boundary information provided by transcriptome annotations. The clustering algorithm uses the minimum distance calculation (Fig. 2) where any HSP within a user-defined distance threshold of an existing group is merged into that putative gene unit. This approach reduces noise from scattered sequence similarities while maintaining sensitivity for detecting genuine gene signals. Similar HSP clustering approaches have been employed in other comparative genomics pipelines [15, 16], which additionally extend beyond signal detection to attempt gene structure prediction, though with variable success. Given that no universally reliable method for computational gene prediction currently exists, *Novabrowse* restricts its scope to gene signal identification and synteny-based candidate area localization, leaving the method of choice for further downstream investigation up to the user.

For the present analysis, we set the minimum HSP clustering distance threshold to 1,200,000 bp based on the genomic architecture of *P. waltl*. Analysis of the current genome annotation (GCF_031143425.1), containing 53,952 genes, reveals a maximum intron length of 1,186,645 bp. By setting our clustering threshold slightly above this maximum, we ensure high probability of detecting gene signals even in cases where large insertion events or extensive intronic expansions have occurred.

#### Identification of putative *Foxp3* gene signals

Based on the preceding synteny analysis, which identified the top of chromosome 10 as the most probable location for *Foxp3* through flanking gene conservation (Fig. 5B), we restricted our genome search to this chromosome. This targeted approach illustrates a workflow in which synteny analysis first narrows the candidate region, allowing subsequent gene signal detection to be focused on the most informative genomic interval.

Using human *FOXP3* as the query sequence, *Novabrowse*’s genome search identified four putative gene signals on *P. waltl* chromosome 10 (Fig. 8A). Two signals, *Gene_2* and *Gene_4* (coordinates 292,144,642– 292,145,978), exhibited the highest coverage (19.5% and 18.4% respectively) but mapped within the bounds of the annotated *Parn* (*poly(A)-specific ribonuclease)* gene (coordinates 292,083,872–292,774,409). The 3’ region of *P. waltl Parn* appears to share similarity with a section in human *FOXP3*, indicating that *Gene_2* and *Gene_4* signals represent matches to an existing gene rather than to a novel *Foxp3* signal. The remaining two signals, *Gene_1* (coordinates 3,749,056–3,749,178) and *Gene_3* (same coordinates), mapped within the syntenic region of interest at the top of chromosome 10, showing coverage of 5.4% and 5.1% respectively and aligning to the middle section of human *FOXP3*. Despite the low coverage of *Gene_1* and *Gene_3*, the convergence of these signals within the syntenic block at the top of chromosome 10 strengthens the case for a potentially unannotated *Foxp3* residing in this region.

**Figure 8.**
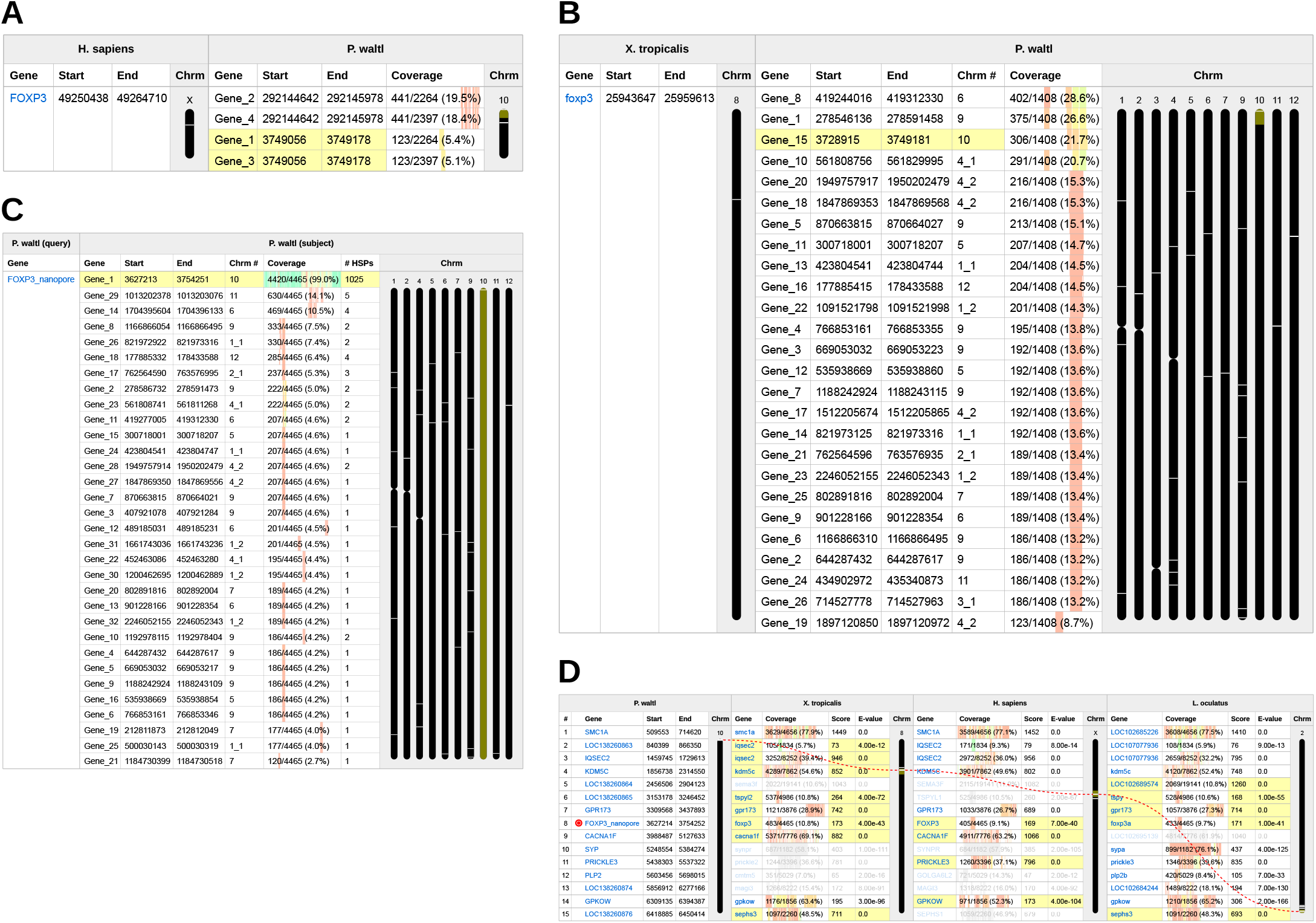
*Foxp3* gene signal detection, validation, and synteny confirmation in *P. waltl* using *Novabrowse*. **(A)** tBLASTx search results of human *FOXP3* transcript against *P. waltl* chromosome 10. **(B)** tBLASTx search results of *X. tropicalis foxp3* transcript against *P. waltl* genome. **(C)** tBLASTx search results of a Nanopore-sequenced *Foxp3* transcript against the *P. waltl* genome, used to validate the gene signals identified in (A) and (B). The “# HSPs” column shows the number of High-scoring Segment Pairs consolidated into each putative gene signal. No chromosome is assigned to *FOXP3_nanopore* (query) as it represents raw Nanopore transcript sequence data not mapped to the genome assembly. **(D)** Synteny analysis of *P. waltl* genes flanking the discovered *FOXP3_nanopore* coordinates (seven genes upstream, seven genes downstream), showing chromosomal positions and syntenic relationships compared across four species: Iberian ribbed newt (*P. waltl*), African clawed frog (*X. tropicalis*), human (*H. sapiens*), and spotted gar (*L. oculatus*). The analysis was performed using tBLASTx searches with *P. waltl* gene sequences as queries against each target species’ transcriptome. *FOXP3_nanopore* was added to the query set using the custom sequence feature. The red dashed ribbon indicates *FOXP3_nanopore*. “Start” and “End” columns indicate chromosomal positions throughout. Chromosome heights are normalized to equal heights. See Fig. 3 for further layout interpretation.

To validate these findings using a phylogenetically closer reference, we repeated the analysis with *X. tropicalis foxp3*, the closest amphibian species with annotated *foxp3* in NCBI databases. Similar to human analysis, the *X. tropicalis foxp3* search against the *P. waltl* transcriptome produced comparable results (Supplemental Fig. 2), and synteny analysis confirmed that the *X. tropicalis foxp3* locus shows close syntenic conservation with the top of *P. waltl* chromosome 10 (Supplemental Fig. 3).

In contrast to the targeted human query, the *X. tropicalis foxp3* search was performed against all *P. waltl* chromosomes (Fig. 8B). *Gene_15* (coordinates 3,728,915–3,749,181) was identified at the top of *P. waltl* chromosome 10, with chromosomal coordinates closely matching those of *Gene_1* and *Gene_3* from the human search. Two additional matches exhibited higher coverage values, however, these mapped to the *Foxp4* genomic region on chromosome 6 (*Gene_8*, 28.6%; coordinates 419,244,016–419,312,330) and the *Foxp1* genomic region on chromosome 9 (*Gene_1*, 26.6%; coordinates 278,546,136–278,591,458). Notably, most matches, including the aforementioned three highest-scoring (i.e., *Gene_8, Gene_1*, and *Gene_15*), showed coverage concentrated in the same region of the *X. tropicalis foxp3* query, located at approximately three-quarters of its length. This shared coverage pattern likely reflects a conserved Forkhead box family domain common to multiple *Foxp* paralogs.

Reciprocal syntenic analysis showed that the region where *Gene_1*/*Gene_3* (human search) and *Gene_15* (*X. tropicalis* search) map, at the top of chromosome 10, is flanked upstream by *Gpr173* and downstream by *Cacna1f*. These same genes flank *Foxp3* orthologs across other vertebrate species: in *X. tropicalis, gpr173* is immediately downstream of *foxp3* and *cacna1f* is two genes upstream (Supplemental Fig. 3); in humans, *CACNA1F* is three genes upstream of *FOXP3* (Fig. 5B).

None of the genomic regions where *Gene_1*/*Gene_3* (human search) and *Gene_15* (*X. tropicalis* search) map have any annotated genes in the current *P. waltl* genome assembly, nor do the regions where the higher-coverage matches *Gene_2* and *Gene_4* (from the human search) align. However, the syntenic conservation combined with the independent identification of putative gene signals at overlapping coordinates by both tBLASTx searches indicates that the gene signal at the top of *P. waltl* chromosome 10 could represent a missing annotation of the *Foxp3* locus. Confirming gene presence at this stage remains beyond the reach of computational methods alone, as no existing tool can reliably predict whether a true gene resides at an identified signal. Resolving such cases ultimately requires experimental evidence, for example, targeted transcriptomic sequencing of tissues where the gene is expected to be expressed.

#### Identification of putative *Aire* gene signals

Having identified putative *Foxp3* gene signals within the expected syntenic region, we next applied the same genome-based approach to investigate the second unannotated immune gene, *Aire*. Using human *AIRE* as the query sequence, *Novabrowse*’s genome search identified six putative gene signals (Fig. 9A). The highest-coverage match *Gene_1* (24.4% coverage) localized to chromosome 11 at coordinates 504,074,795– 504,155,916, aligning primarily to *AIRE’s* 5’ region and central regions. Five additional signals with lower coverage (3.7-6.2%) scattered across chromosomes 3, 6, 7, and 12.

**Figure 9.**
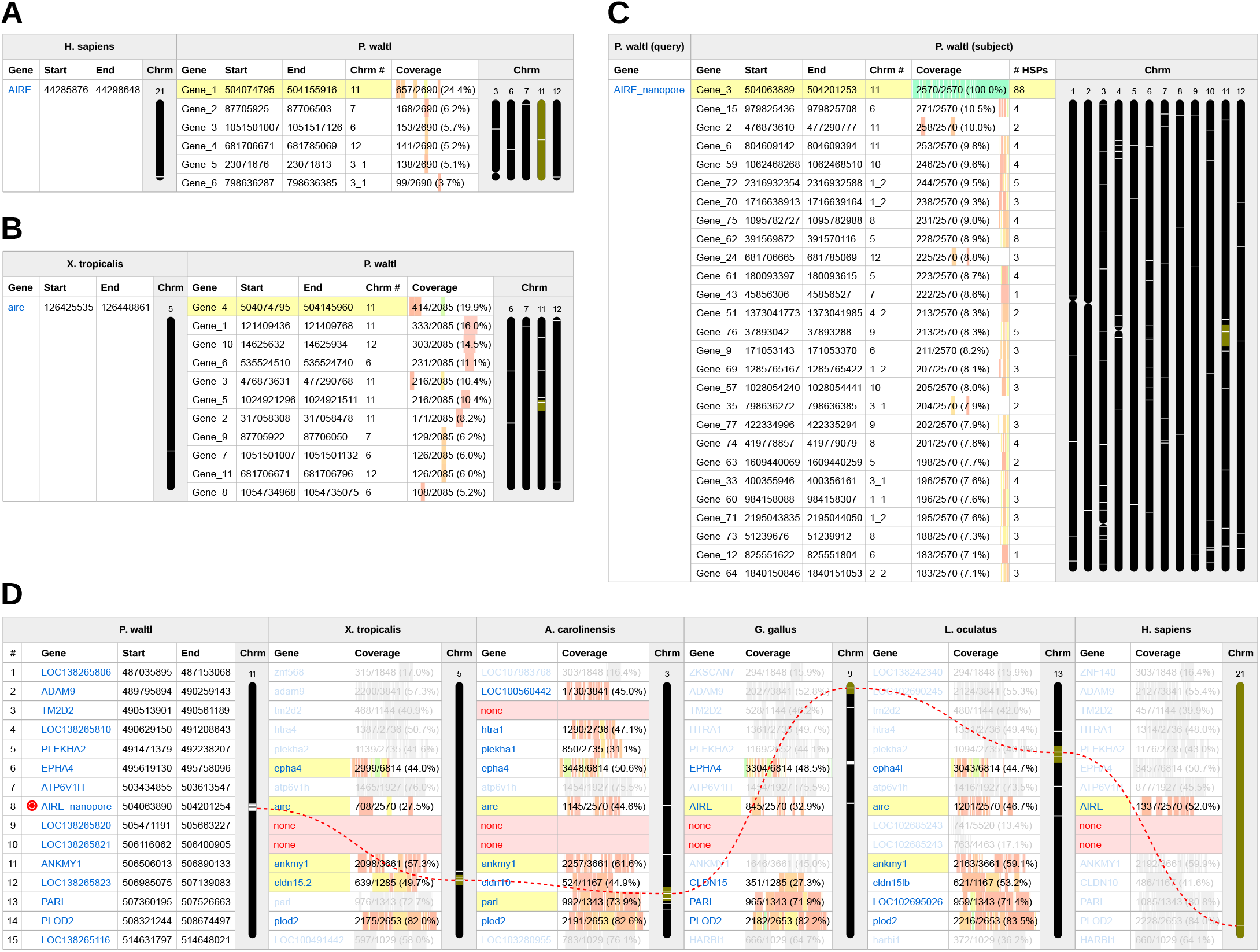
*Aire* gene signal detection, validation, and synteny confirmation in *P. waltl* using *Novabrowse*. **(A)** tBLASTx search results of human *AIRE* transcript against the *P. waltl* genome. **(B)** tBLASTx search results of *X. tropicalis aire* transcript against the *P. waltl* genome. **(C)** tBLASTx search results of a Nanopore-sequenced *Aire* transcript against the *P. waltl* genome, used to validate the gene signals identified in (A) and (B). The “# HSPs” column shows the number of High-scoring Segment Pairs consolidated into each putative gene signal. No chromosome is assigned to *AIRE_nanopore* (query) as it represents raw Nanopore transcript sequence data not mapped to the genome assembly. On the chromosome visualization, all matches are shown, while the table displays only matches with >7% coverage. See Supplemental Fig. 4 for complete results. **(D)** Synteny analysis of *P. waltl* genes flanking the discovered *AIRE_nanopore* coordinates (seven genes upstream, seven genes downstream), showing chromosomal positions and syntenic relationships compared across six species: Iberian ribbed newt (*P. waltl*), African clawed frog (*X. tropicalis*), lizard (*A. carolinensis*), chicken (*G. gallus*), spotted gar (*L. oculatus*), and human (*H. sapiens*). The analysis was performed using tBLASTx searches with *P. waltl* gene sequences as queries against each target species’ transcriptome. *AIRE_nanopore* was added to the query set using the custom sequence feature. The red dashed ribbon indicates *AIRE_nanopore*. “Start” and “End” columns indicate chromosomal positions throughout. Chromosome heights are normalized to equal heights. See Fig. 3 for further layout interpretation.

The *X. tropicalis aire* search against *P. waltl* genome yielded eleven signals (Fig. 9B), with *Gene_4* producing the highest-coverage match on chromosome 11 at coordinates (504,074,795–504,145,960) that shares an identical start position with *Gene_1* from the human search but differs at the end coordinates. Interestingly, despite *X. tropicalis* being phylogenetically closer to *P. waltl* than humans, *Gene_4* shows lower coverage (19.9%) than *Gene_1* from the human search (24.4%). This difference is particularly notable given the shorter *X. tropicalis aire* query sequence (2,085 bp versus 2,690 bp), suggesting that *AIRE* sequence conservation may not have been uniformly preserved across amphibian lineages.

The regions where *Gene_4* (*X. tropicalis* search) and *Gene_1* (*human* search) map fall within the previously identified syntenic block on chromosome 11 (Fig. 7). Similar to the *Foxp3* findings, no genes are currently annotated in this genomic region. However, unlike the *Foxp3* analysis where synteny clearly delineated a single candidate region, the *Aire* signal showed greater dispersion reflecting the expansion of that region in *P. waltl* compared to *X. tropicalis*. The closest syntenic match is *Ankmy1*, which is located two genes downstream from where these gene signals map in *P. waltl*, while in *X. tropicalis, ankmy1* is located 21 genes downstream from *aire*.

Despite this weaker syntenic context, the overlapping coordinates from both searches still suggest that the area of *Gene_4* (*X. tropicalis* search) and *Gene_1* (human search) represents the putative *Aire* locus. As with *Foxp3*, experimental validation is necessary to determine whether a functional gene is present at this location.

### Gene Signal Confirmation

The preceding analyses identified putative *Foxp3* and *Aire* gene signals at specific genomic coordinates in *P. waltl* through distance-based HSP clustering. We hypothesized that *Novabrowse* predictions indicate these genes are present, but that a lack of RNA sequencing data has precluded their annotation. To validate these predictions and assess the robustness of *Novabrowse*’s approach, we performed RNA sequencing followed by comparative genomic localization using both conventional methods and *Novabrowse*.

#### Nanopore sequencing and independent genomic localization

Given that one probable explanation for the lack of *Foxp3* and *Aire* annotations is the absence of transcriptomic data from tissues where these genes are primarily expressed, we targeted the thymus for sequencing. The thymus is a small primary lymphoid organ central to adaptive immunity, where both *Foxp3* and *Aire* are known to be expressed in other vertebrates, increasing the likelihood of capturing these transcripts. The sequencing yielded full-length transcripts that were processed through the Oxford Nanopore wf-transcriptomes pipeline, generating a merged transcriptome that included transcripts mapping to unannotated genomic regions (class code=u).

To identify candidate *Foxp3* and *Aire* transcripts among the unannotated sequences, we predicted open reading frames using TransDecoder and screened the resulting peptides via BLASTp against the UniPro - tKB/Swiss-Prot database. This analysis identified two candidates: a 4,465 nt transcript with highest similarity to human *FOXP3* (E-value: 8.58×10^−61^; 38.52% identity), and a 2,570 nt transcript with highest similarity to human *AIRE* (E-value: 4.47×10^−128^; 42.56% identity). To determine the genomic placement of these candidates, the transcript nucleotide sequences were aligned to the *P. waltl* genome assembly using megablast, which localized the *Foxp3* candidate to chromosome 10 (coordinates 3,627,213–3,752,105) and the *Aire* candidate to chromosome 11 (coordinates 504,063,889–504,201,253).

#### *Novabrowse*-based genomic localization

Using *Novabrowse*’s custom query feature, the Nanopore-derived transcripts (designated FOXP3_nanopore and AIRE_nanopore) were matched against the *P. waltl* genome assembly using tBLASTx search.

For *Foxp3*, the best match (*Gene_1*) was identified on chromosome 10 at coordinates 3,627,213–3,754,251 with 99% coverage (4,420/4,465 bp) consolidated from 1,025 HSPs (Fig. 8C). This near-complete coverage demonstrates that the distance-based clustering algorithm successfully grouped over a thousand individual alignment segments into a single coherent gene unit spanning approximately 127 kb of genomic sequence. The 1% coverage gap reflects the inherent behavior of BLAST’s heuristic algorithm, which uses seeding and extension-dropoff mechanisms that may not capture every nucleotide position, particularly at HSP boundaries [5,19].

The best match for *Aire* (*Gene_3*) was mapped to chromosome 11 at coordinates 504,063,889–504,201,253 with 100% coverage (2,570/2,570 bp) from 88 HSPs (Fig. 9C). The complete coverage indicates that the entire transcript sequence aligned to a ∼137 kb genomic region, with all HSPs consolidated into a single gene unit.

The coordinates obtained through *Novabrowse* closely match those from megablast, with both methods independently converging on the same chromosomal regions. The agreement between nucleotide-based (megablast) and translated (tBLASTx) approaches further strengthens confidence in the identified loci.

#### Synteny confirmation of orthology

To confirm that the identified transcripts represent true *Foxp3* and *Aire* orthologs rather than related paralogs, we performed synteny analysis using the Nanopore-derived sequences and their genomic flanking genes as queries.

For *Foxp3*, analysis of seven upstream and seven downstream genes flanking the *FOXP3_nanopore* locus revealed strong syntenic conservation across *X. tropicalis, H. sapiens*, and *L. oculatus* (Fig. 8D), confirming orthology through conserved genomic context.

For *Aire*, synteny analysis of flanking genes around *AIRE_nanopore* showed conservation across *X. tropicalis, A. carolinensis, G. gallus, L. oculatus*, and *H. sapiens* (Fig. 9D). While the genomic region exhibits expansion in *P. waltl* relative to other species, consistent with earlier observations (Fig. 7), the overall syntenic relationships support orthology assignment.

#### Concordance with gene signal predictions

The genomic coordinates identified through Nanopore transcript mapping encompass the gene signals detected in our earlier searches using human and *X. tropicalis* queries. For *Foxp3*, the signals identified using human *FOXP3* (*Gene_1* and *Gene_3* at coordinates 3,749,056–3,749,178; Fig. 8A) and *X. tropicalis foxp3* (Gene_15 at 3,728,915–3,749,181; Fig. 8B) fall entirely within the Nanopore-confirmed region of 3,627,213– 3,754,251 (Fig. 8C). Similarly, for *Aire*, both the human-derived signal (*Gene_1* at 504,074,795–504,155,916; Fig. 9A) and the *X. tropicalis*-derived signal (*Gene_4* at 504,074,795–504,145,960; Fig. 9B) map within the Nanopore-confirmed boundaries of 504,063,889–504,201,253 (Fig. 9C).

This concordance validates that *Novabrowse*’s distance-based HSP clustering can pinpoint the location of unannotated genes even when dealing with partial gene signals with modest coverage (5–24%) derived from evolutionarily divergent query species. Despite the approximately 350 and 290 million years separating human and *X. tropicalis* from *P. waltl* respectively [20], the conserved regions detected by both queries were sufficient to identify the correct genomic loci. Notably, sequence conservation at individual loci did not always scale predictably with phylogenetic distance, as the human *AIRE* search yielded higher coverage (24.4%) than the phylogenetically closer *X. tropicalis* search (19.9%).

### High Resolution Chromosomal Rearrangement and Gene Loss Identification

The preceding analyses demonstrated that *Novabrowse* can identify gene signals for loci that appear absent from current annotations but are in fact present in the genome. However, an equally important capability is determining when a gene is genuinely missing rather than unannotated. The retinoblastoma gene family comprises three tumor suppressor genes, *RB1, RBL1*, and *RBL2*, that play essential roles in cell cycle regulation, differentiation, and development across vertebrates [21, 22, 23]. While all three genes are well-conserved across vertebrates, *Rbl1* has not yet been annotated in *P. waltl*. As with *Foxp3* and *Aire*, this raises the question of whether the gene has been lost during newt evolution or simply remains unannotated due to sequence divergence or assembly gaps.

Notably, *Rbl1* is annotated in the closely related salamander species *Ambystoma mexicanum* (axolotl), providing a suitable reference for investigating its status in *P. waltl*. We performed synteny analysis using *A. mexicanum Rbl1* and its 14 flanking genes as queries, comparing chromosomal arrangements across six vertebrate species (Fig. 10A). The analysis revealed strong conservation of the *Rbl1* genomic neighborhood in most species examined. Key syntenic markers including *Samhd1, Mtcl2, Chd6*, and *Ralgapb* consistently localized near *Rbl1* orthologs in *A. mexicanum, X. tropicalis, A. carolinensis, G. gallus, H. sapiens*, and *L. oculatus*. However, in *P. waltl*, while *Mtcl2, Samhd1*, and *Ralgapb* mapped to chromosome 7 in the expected syntenic configuration, both *Rbl1* and *Chd6* were absent from this region. Instead, the highest-scoring matches for axolotl *Rbl1* corresponded to *Rbl2* on *P. waltl* chromosome 12, while axolotl *Chd6* matched to *Chd7* on chromosome 2_2 (Fig. 10A).

**Figure 10.**
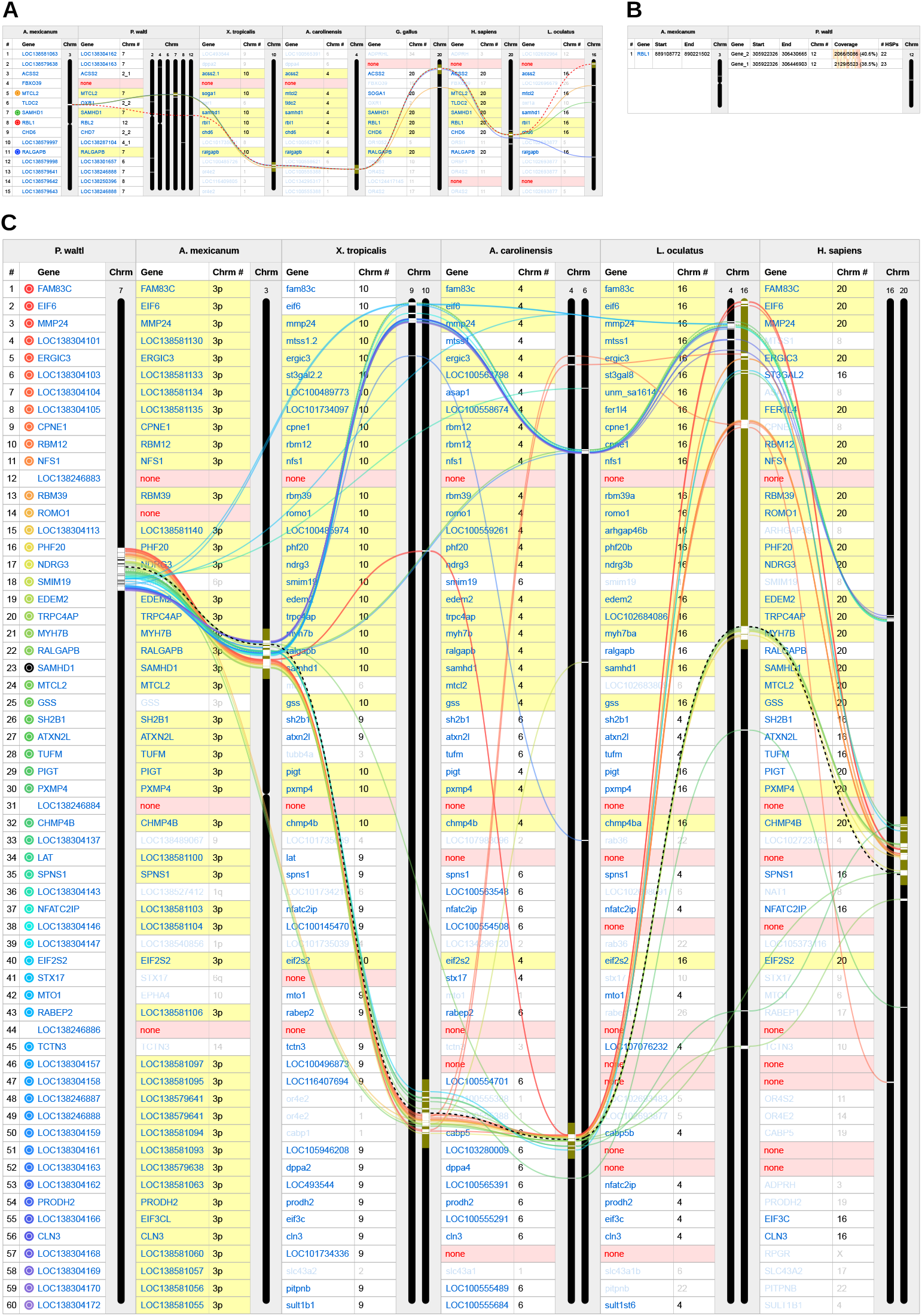
Cross-species comparison of the *Rbl1* genomic neighborhood, *Rbl1* gene signal detection, and chromosomal rearrangement analysis of the expected *Rbl1* locus in *P. waltl* using *Novabrowse*. **(A)** Cross-species synteny comparison of axolotl (*A. mexicanum*) *Rbl1* and 14 flanking genes (seven upstream, seven downstream), showing chromosomal positions and syntenic relationships compared across six species: Iberian ribbed newt (*P. waltl*), African clawed frog (*X. tropicalis*), lizard (*A. carolinensis*), chicken (*G. gallus*), human (*H. sapiens*), and spotted gar (*L. oculatus*). The analysis was performed using tBLASTx searches with *A. mexicanum* gene sequences as queries against each target species’ transcriptome. Query genes are listed in the leftmost column. Colored ribbons indicate enabled gene tracking: *Rbl1* (red, dashed), *Mtcl2* (orange, solid), *Samhd1* (green), and *Ralgapb* (blue). **(B)** tBLASTx search results of *A. mexicanum Rbl1* transcript against the *P. waltl* genome. The “# HSPs” column shows the number of High-scoring Segment Pairs consolidated into each putative gene signal. **(C)** Synteny analysis of 60 *P. waltl* genes from the genomic region where *Rbl1* would be expected to reside, showing chromosomal positions and syntenic relationships compared across five species: axolotl (*A. mexicanum*), African clawed frog (*X. tropicalis*), lizard (*A. carolinensis*), human (*H. sapiens*), and spotted gar (*L. oculatus*). The analysis was performed using tBLASTx searches with *P. waltl* gene sequences as queries against each target species’ transcriptome. Query genes are listed in the leftmost column. The black dashed ribbon indicates *Samhd1*. Yellow backgrounds indicate genes highlighted through coordinate-based area filtering, marking the syntenic block corresponding to the *Rbl1* locus in those species. For *L. oculatus*, the highlight was expanded to include both the *rbl1* locus at the top of chromosome 16 and *samhd1*, showing that despite genomic expansion of this locus in spotted gar, most syntenic genes have remained on the same chromosome. “Start” and “End” columns indicate chromosomal positions throughout. Chromosome heights are normalized to equal heights. See Fig. 3 for further layout interpretation.

Gene signal detection searches provided further evidence for *Rbl1* absence. When *A. mexicanum Rbl1* was queried against the *P. waltl* genome, *Novabrowse* identified only two putative gene signals, both localizing to chromosome 12 with 40.6% and 38.5% coverage respectively (Fig. 10B). Examination of these genomic coordinates confirmed they corresponded to the annotated *Rbl2* locus rather than a distinct *Rbl1* ortholog. Searches using human *RBL1* as the query sequence yielded 11 putative gene signals (Supplemental Fig. 5). Similar to the axolotl query, ten of these signals mapped to chromosome 12, all within the same genomic region where *Rbl2* resides, with the highest coverage matches reaching 71.2% and 62.3%. An additional match on chromosome 8 with 6.3% coverage corresponded to the *P. waltl Rb1* locus. These results reflect the conserved functional domains shared among retinoblastoma gene family members rather than the presence of a true *Rbl1* ortholog.

To understand the genomic context of this apparent gene loss, we used *Novabrowse* to perform high-resolution synteny analysis of 60 genes spanning the expected *Rbl1* locus on *P. waltl* chromosome 7 (Fig. 10C). This analysis revealed a striking pattern of chromosomal rearrangement. In *A. mexicanum*, the majority of these genes localize to a defined region in chromosome 3p, where *Rbl1* also resides (Fig. 10A), indicating that this genomic region has been largely preserved between the two salamander species. However, when compared to a fellow amphibian *X. tropicalis* or to more distantly related vertebrates including *A. carolinensis, H. sapiens*, and *L. oculatus*, the syntenic block splits across two distinct chromosomal locations. Genes upstream of *Mtcl2 and Samhd1* (including *Ralgapb*) map to one chromosome, while genes beginning approximately three positions downstream of *Samhd1*, starting with *Sh2b1*, predominantly map to a distinct locus on a different chromosome in these species. Notably, the spotted gar *L. oculatus* also exhibits this two-chromosome arrangement, suggesting it represents the ancestral vertebrate state. This pattern indicates that the salamander lineage experienced a chromosomal fusion or translocation event that joined two ancestrally separate genomic regions.

Since both salamander species share the fused chromosomal arrangement, the rearrangement itself likely occurred in their common ancestor and initially preserved *Rbl1*, as evidenced by its retention in *A. mexicanum*. The loss of *Rbl1* from *P. waltl*, likely along with *Chd6* which also shows no ortholog in the expected syntenic position, therefore appears to represent a subsequent, lineage-specific deletion. The complete absence of any sequence with significant homology to *Rbl1* across both transcriptome and genome-wide searches supports genuine gene loss rather than an assembly gap or annotation artifact.

## Discussion

Gene annotation remains fundamentally constrained by an incomplete mechanistic understanding of how genes are realized as functional units. While transcriptomic data can often pinpoint where genes reside in the genome, determining their specific function based on sequence alone remains a challenge. In the absence of a complete theoretical framework, comparison based on empirical data from related species remains our most reliable approach for identifying gene presence and inferring their function. Current annotation pipelines rely heavily on transcriptomic evidence, however when such data is incomplete, as seen with *Foxp3* and *Aire*, the resulting annotations, or lack thereof, can be misleading.

A common first step in such comparative validation is sequence-similarity searching with BLAST, which can rapidly identify candidate homologs and provide an initial ranking of matches based on alignment statistics. BLAST outputs are highly effective for answering whether homologous sequences exist, but they are less well suited to distinguishing true orthologs from paralogs, assessing the genomic context of matches, or integrating evidence from multiple species simultaneously. Existing tools address parts of this problem but operate at either genome-wide or single-query scales, leaving a gap at the intermediate resolution where annotation ambiguities are actually resolved.

We developed *Novabrowse* to fill this gap. By integrating BLAST homology searches with interactive synteny visualization, it enables researchers investigating individual genes or broader genomic regions to screen candidate homologs, assess their chromosomal distribution, and evaluate syntenic relationships across multiple reference species within a single integrated pipeline.

As the feature comparison illustrates (Table 1), while *Novabrowse* introduces novel features such as coordinate-based region search and isoform-aware hit consolidation, its primary strength lies in unifying capabilities that have traditionally required researchers to move between multiple platforms. By bringing these steps into a single pipeline, *Novabrowse* reduces the practical bottleneck in comparative genomics workflows, which, for many researchers, is often not the alignment speed but rather the manual effort of integrating outputs across disconnected platforms.

We demonstrated these capabilities through analysis of *Foxp3* and *Aire* in *P. waltl*. By integrating BLAST results with interactive synteny visualization, the tool enabled us to: rapidly screen 38 *FOXP3* and nine *AIRE* homologous matches across their chromosomal distribution; visualize alignment coverage to assess match quality and completeness at a glance; examine syntenic relationships across multiple species simultaneously at flexible resolution scales (from 14 to 60 flanking genes); seamlessly adapt our analysis strategy by switching reference species when human synteny proved inadequate; track syntenic blocks even through extreme genomic expansions and rearrangements, and focus analysis on specific chromosomal regions through coordinate-based filtering. This integrated approach, combining sequence similarity, genomic context, and coverage metrics in a single interface, revealed patterns that would be difficult to detect using traditional BLAST output or any currently existing tools alone.

The Nanopore-based validation of *Foxp3* and *Aire* putative gene signals additionally addressed a methodological question: whether *Novabrowse*’s distance-based HSP clustering can reliably identify genes in repeat-rich, large genomes. The *P. waltl* genome presents a challenging test case, with its 20.3 Gb size and extensive repetitive content. The 99–100% coverage achieved when mapping species-specific transcripts back to the genome confirms that the 1,200,000 bp clustering threshold (derived from maximum annotated intron length) effectively consolidates related HSPs without over-grouping unrelated sequences.

The analysis of *Rbl1* demonstrated a complementary but distinct outcome of the same analytical framework. High-resolution synteny analysis revealed that the genomic region corresponding to the ancestral *Rbl1* locus underwent a chromosomal fusion or translocation event in the salamander lineage, joining two ancestrally separate genomic regions. Subsequently, *Rbl1* itself, likely along with *Chd6*, was lost from *P. waltl* despite retention of flanking genes. Notably, both *Rbl1* and *Chd6* remain present in *A. mexicanum*, indicating that the loss was specific to the *P. waltl* lineage. Genome-wide searches using both *A. mexicanum Rbl1* and human *RBL1* as queries further supported this conclusion, as all high-scoring matches corresponded to other retinoblastoma family members rather than a distinct *Rbl1* ortholog (Fig. 10B; Supplemental Fig. 5). These findings highlight Novabrowse’s ability to resolve complex evolutionary history questions. While the cause of this lineage-specific deletion remains unclear, the synteny-based framework enabled its detection and contextualization within the broader chromosomal landscape.

*Novabrowse* was designed with a specific analytical niche in mind, and its current implementation reflects certain trade-offs. The tool’s reliance on NCBI E-utilities for query sequence retrieval means that species or genes absent from public databases require manual input of custom sequences, though this functionality is supported. While independent of the tool itself, the choice of search algorithm introduces computational considerations. This is particularly relevant for tBLASTx, which provides the sensitivity needed to detect homologs across large evolutionary distances but requires substantially more processing time than BLASTn or tBLASTn searches. The interactive HTML output also imposes practical constraints on analysis scale. Displaying approximately 100 query genes across six subject species approaches the upper limit that modern browsers can render without significant performance degradation, which may necessitate splitting larger analyses into smaller batches.

Building on the current implementation, several enhancements would further expand the tool’s utility. Integration of alternative alignment algorithms could reduce search times when matching multiple query genes against multiple subject species. DIAMOND [24] offers substantially faster protein-based search, though at the cost of reduced sensitivity relative to BLAST. MMseqs2 [25] provides even greater speed improvements and would be particularly valuable for rapid preliminary surveys, with users able to follow up promising hits using more sensitive BLAST searches if necessary. Additionally, expanding annotation file support to include GFF/GFF3 formats alongside the current GTF requirement would increase compatibility with a wider range of genome databases and annotation sources.

*Novabrowse* is released under the MIT license and is freely available at https://github.com/RegenImm-Lab/Novabrowse with documentation and example datasets to facilitate adoption. We envision this tool being particularly valuable for researchers working with non-model organisms, where annotation quality often lags behind assembly quality, and for comparative genomics studies that require systematic evaluation of gene presence, absence, or orthology across multiple species. The modular design allows integration into existing genomics workflows, with the pipeline being implemented entirely in Jupyter notebook format, making it straightforward to modify and adapt the code to specific needs. We also provide a containerized version via Docker, which is also compatible with Apptainer, allowing a ready-to-use deployment option that simplifies installation and use.

## Conclusions

*Novabrowse* provides users with an integrated platform for BLAST results interpretation that combines homology search with synteny analysis at a resolution scale not well served by existing tools. Through case studies in *P. waltl*, we demonstrated its utility for ortholog detection, gene signal discovery in unannotated regions, and chromosomal rearrangement analysis. We also showed that even low-coverage matches from divergent query species can accurately pinpoint gene loci when sequence conservation is limited to specific functional domains.

The challenges encountered in identifying *Foxp3* and *Aire* in *P. waltl* highlight a broader concern for genome annotation. As an emerging model organism, *P. waltl* can be bred in laboratory settings, providing relatively abundant tissue access for transcriptomics. Yet despite these resources, both genes remained unannotated until targeted thymus sequencing was performed.

As genome sequencing increasingly extends to species where tissue sampling is limited, such annotation gaps are likely to become even more prevalent. Tools, like *Novabrowse*, that enable evidence-based evaluation of gene presence independent of existing annotations will therefore become increasingly important.

## Supporting information

Pwaltl_Aire_and_Foxp3_sequences

## Data Availability

Oxford Nanopore cDNA sequencing data are available at NCBI under BioProject accession PRJNA1443387 as FASTQ files. *Novabrowse* source configurations used to generate all analyses presented in this study along with corresponding output HTML files are available in the project repository in the manuscript_analyses folder at https://github.com/RegenImm-Lab/Novabrowse.

## Acknowledgements

This work was supported by funding from the Swedish Research Council (2020-01486), Cancerfonden (23 3047 Pj), Knut and Alice Wallenberg Foundation, and Crafoord Foundation (#20230825) to N.D.L.. We are grateful to the StemTherapy Strategic Research Area at Lund University for their support. Computations and data storage were enabled by resources provided by LUNARC, The Centre for Scientific and Technical Computing at Lund University. We thank Lisbeth Verk for helpful advice and feedback. We also acknowledge the National Bioinformatics Infrastructure Sweden (NBIS) at SciLifeLab, who were commissioned to develop the Docker container for *Novabrowse*.

