## Supplemental Figures for "*Novabrowse:* A Tool for High-Resolution Synteny Analysis, Ortholog Detection, and Gene Signal Discovery"

| X. tropicalis | | P. waltl | | | | | |
|---|---|---|---|---|---|---|---|
| **Gene** | **Chrm** | **Gene** | **Chrm #** | **Coverage** | **Score** | **E-value** | **Chrm** |
| aire | 5 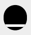 | LOC138265795 | 11 | 357/2085 (17.1%) | 93 | 2.00e-26 | 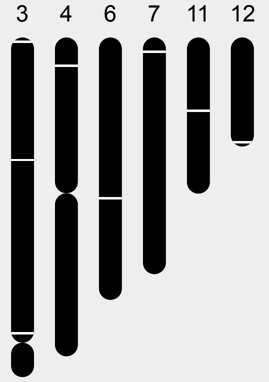  |
| | | TRIM24 | 4_1 | 138/2085 (6.6%) | 84 | 3.00e-15 | |
| | | CHD4 | 7 | 141/2085 (6.8%) | 83 | 8.00e-15 | |
| | | CHD5 | 6 | 138/2085 (6.6%) | 81 | 2.00e-14 | |
| | | TRIM66 | 3_1 | 144/2085 (6.9%) | 79 | 1.00e-13 | |
| | | CHD3 | 12 | 138/2085 (6.6%) | 79 | 1.00e-13 | |
| | | PHF21A | 3_1 | 315/2085 (15.1%) | 75 | 5.00e-13 | |
| | | PHF12 | 3_1 | 234/2085 (11.2%) | 64 | 2.00e-15 | |

**Supplemental Figure 1.** ***Aire*** **ortholog search results in** ***P. waltl*** **using** ***X. tropicalis aire*** **as query.** tBLASTx search results against the *P. waltl* transcriptome visualized with *Novabrowse* showing *X. tropicalis aire* and its homologous matches in *P. waltl* (see Figure 3 for layout interpretation). Chromosome sizes are scaled proportionally to their actual lengths.

| X. tropicalis | | | | P. waltl | | | | | | | |
|---|---|---|---|---|---|---|---|---|---|---|---|
| Gene | Start | End | Chrm | Gene | Start | End | Chrm # | Coverage | Score | E-value | Chrm |
| foxp3 | 25943647 | 25959613 | 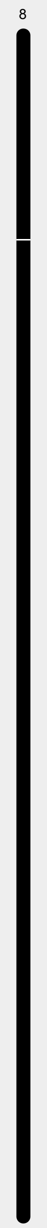 | FOXP1 | 277849797 | 278617590 | 9 | 507/1408 (36.0%) | 195 | 3.00e-58 | 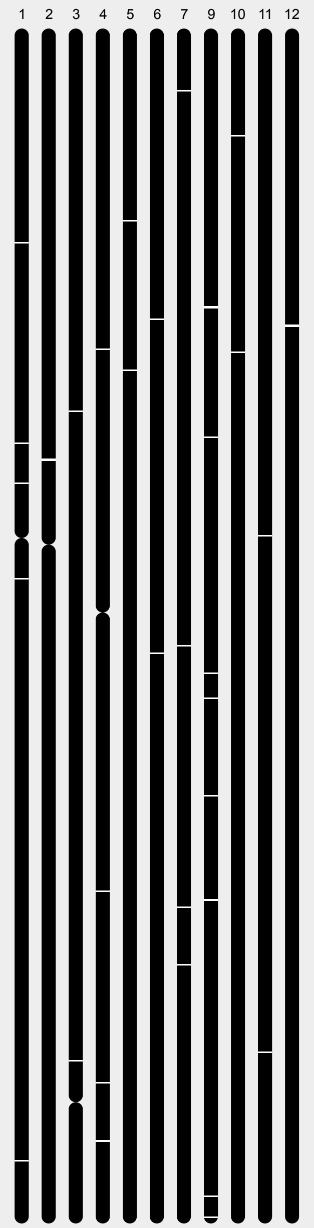 |
| | | | | FOXP4 | 419038116 | 419327081 | 6 | 465/1408 (33.0%) | 186 | 2.00e-53 | |
| | | | | FOXP2 | 561752083 | 562013147 | 4_1 | 468/1408 (33.2%) | 177 | 2.00e-51 | |
| | | | | LOC138259942 | 766851390 | 766865484 | 9 | 195/1408 (13.8%) | 90 | 2.00e-17 | |
| | | | | FOXJ1 | 1143257857 | 1143273636 | 7 | 225/1408 (16.0%) | 88 | 1.00e-16 | |
| | | | | FOXA3 | 669047961 | 669183575 | 9 | 192/1408 (13.6%) | 88 | 2.00e-16 | |
| | | | | FOXA2 | 535937538 | 535943925 | 5 | 192/1408 (13.6%) | 86 | 4.00e-16 | |
| | | | | LOC138261469 | 293399302 | 293417124 | 10 | 312/1408 (22.2%) | 85 | 7.00e-17 | |
| | | | | FOXA1 | 1188237926 | 1188338696 | 9 | 192/1408 (13.6%) | 85 | 1.00e-15 | |
| | | | | FOXD1 | 423803004 | 423806419 | 1_1 | 204/1408 (14.5%) | 84 | 2.00e-15 | |
| | | | | LOC138293472 | 1949755843 | 1949759072 | 4_2 | 216/1408 (15.3%) | 84 | 3.00e-15 | |
| | | | | FOXC2 | 178338656 | 178342146 | 12 | 189/1408 (13.4%) | 84 | 3.00e-15 | |
| | | | | FOXG1 | 1166865623 | 1166920147 | 9 | 186/1408 (13.2%) | 84 | 3.00e-15 | |
| | | | | FOXC1 | 763576084 | 763578164 | 2_1 | 189/1408 (13.4%) | 83 | 5.00e-15 | |
| | | | | FOXN3 | 870429141 | 871077790 | 9 | 195/1408 (13.8%) | 82 | 7.00e-15 | |
| | | | | FOXF1 | 177884128 | 177902644 | 12 | 186/1408 (13.2%) | 82 | 1.00e-14 | |
| | | | | FOXJ2 | 79961724 | 80152668 | 7 | 186/1408 (13.2%) | 81 | 2.00e-14 | |
| | | | | FOXN2 | 300693584 | 300805818 | 5 | 207/1408 (14.7%) | 81 | 2.00e-14 | |
| | | | | LOC138293902 | 1512205551 | 1512206351 | 4_2 | 192/1408 (13.6%) | 81 | 2.00e-14 | |
| | | | | FOXL1 | 178433168 | 178435723 | 12 | 201/1408 (14.3%) | 81 | 2.00e-14 | |
| | | | | LOC138284875 | 821972707 | 821974482 | 1_1 | 192/1408 (13.6%) | 80 | 3.00e-14 | |
| | | | | FOXD3 | 1847868120 | 1847869993 | 4_2 | 216/1408 (15.3%) | 80 | 4.00e-14 | |
| | | | | LOC138294231 | 1950200250 | 1950202797 | 4_2 | 201/1408 (14.3%) | 79 | 5.00e-14 | |
| | | | | FOXF2 | 762562737 | 762588324 | 2_1 | 186/1408 (13.2%) | 79 | 9.00e-14 | |
| | | | | FOXI3 | 2246048086 | 2246052771 | 1_2 | 189/1408 (13.4%) | 78 | 1.00e-13 | |
| | | | | FOXL2 | 435338099 | 435341332 | 11 | 186/1408 (13.2%) | 78 | 2.00e-13 | |
| | | | | FOXL3 | 96738634 | 96757616 | 10 | 246/1408 (17.5%) | 78 | 2.00e-13 | |
| | | | | LOC138258691 | 644285903 | 644287983 | 9 | 186/1408 (13.2%) | 78 | 2.00e-13 | |
| | | | | FOXE1 | 1091518351 | 1091525569 | 1_2 | 201/1408 (14.3%) | 77 | 3.00e-13 | |
| | | | | FOXI1 | 802889264 | 802892441 | 7 | 189/1408 (13.4%) | 76 | 5.00e-13 | |
| | | | | FOXI2 | 901227408 | 901232329 | 6 | 189/1408 (13.4%) | 76 | 6.00e-13 | |
| | | | | FOXB1 | 714522909 | 714530859 | 3_1 | 186/1408 (13.2%) | 76 | 6.00e-13 | |
| | | | | FOXQ1 | 762062105 | 762064987 | 2_1 | 231/1408 (16.4%) | 74 | 2.00e-12 | |
| | | | | FOXN1 | 1927621261 | 1928021333 | 3_1 | 204/1408 (14.5%) | 74 | 2.00e-12 | |
| | | | | FOXN4 | 879085299 | 879137628 | 11 | 270/1408 (19.2%) | 73 | 2.00e-12 | |
| | | | | FOXK2 | 1218268597 | 1218491773 | 7 | 186/1408 (13.2%) | 73 | 3.00e-12 | |
| | | | | FOXB2 | 901729236 | 901731219 | 1_1 | 186/1408 (13.2%) | 73 | 4.00e-12 | |
| | | | | LOC138259414 | 407784970 | 407995297 | 9 | 189/1408 (13.4%) | 73 | 4.00e-12 | |

**Supplemental Figure 2. *Foxp3* ortholog search results in *P. waltl* using *X. tropicalis foxp3* as a query.** tBLASTx search results against the *P. waltl* transcriptome visualized with *Novabrowse*. "Start" and "End" columns indicate chromosomal positions. Chromosome heights are normalized to equal heights. Fig. 3 for table layout interpretation

| X. tropicalis | | | P. waltl | | | H. sapiens | | | M. musculus | | | L. oculatus | | |
|---|---|---|---|---|---|---|---|---|---|---|---|---|---|---|
| # | Gene | Chrm | Gene | Chrm # | Chrm | Gene | Chrm # | Chrm | Gene | Chrm # | Chrm | Gene | Chrm # | Chrm |
| 1 | slc27a4 | 8 | SLC27A4 | 6 | 1 4 6 9 10 | SLC27A4 | 9 | X | Slc27a4 | 2 | X | slc27a4 | 24 | 2 |
| 2 | urm1 | | URM1 | 6 | | URM1 | 9 | | Urm1 | 2 | | urm1 | 24 | |
| 3 | cercam | | CERCAM | 6 | | CERCAM | 9 | | Cercam | 2 | | cercam | 24 | |
| 4 | XB997834 | | CCNO | 1_1 | | CCNO | 5 | | Ccno | 13 | | LOC102684308 | 3 | |
| 5 | LOC101730848 | | CCNO | 1_1 | | CCNO | 5 | | Ccno | 13 | | LOC102684308 | 3 | |
| 6 | cacna1f | | CACNA1F | 10 | | CACNA1F | X | | Cacna1f | X | | cacna1fa | 2 | |
| 7 | ccdc22 | | CCDC22 | 10 | | CCDC22 | X | | Ccdc22 | X | | ccdc22 | 2 | |
| 8 | foxp3 | | FOXP1 | 9 | | FOXP3 | X | | Foxp3 | X | | foxp3a | 2 | |
| 9 | gpr173 | | GPR173 | 10 | | GPR173 | X | | Gpr173 | X | | gpr173 | 2 | |
| 10 | tspyl2 | | LOC138260865 | 10 | | TSPYL4 | 6 | | Tspyl4 | 10 | | tspy | 2 | |
| 11 | sema3fl | | LOC138260864 | 10 | | SEMA3F | 3 | | Sema3f | 9 | | LOC102689574 | 2 | |
| 12 | kdm5c | | KDM5A | 4_1 | | KDM5C | X | | Kdm5c | X | | kdm5c | 2 | |
| 13 | iqsec2 | | IQSEC2 | 10 | | IQSEC2 | X | | Iqsec2 | X | | LOC107077936 | 2 | |
| 14 | fcn2 | | LOC138301222 | 6 | | FCN3 | 1 | | Fcnb | 2 | | LOC138241264 | 9 | |
| 15 | ubl4a | | none | | | UBL4A | X | | Ubl4a | X | | ubl4a | 2 | |

**Supplemental Figure 3. Synteny analysis of the *foxp3* loci across vertebrate species using *X. tropicalis* as the query species.** *Novabrowse* visualization showing chromosomal positions and syntenic relationships of *X. tropicalis foxp3* and its 14 flanking genes (seven upstream, seven downstream) compared across five species: African clawed frog (*X. tropicalis*), Iberian ribbed newt (*P. waltl*), human (*H. sapiens*), mouse (*M. musculus*), and spotted gar (*L. oculatus*). The analysis was performed using tBLASTx searches with *X. tropicalis* gene sequences as queries against each target species' transcriptome, followed by chromosomal filtering to display only chromosomes with *foxp3* matches (excluding *P. waltl* where all chromosomes with matches are shown). The red circle before *foxp3* indicates its ribbon is enabled, which appears as a red dashed line tracking *foxp3*'s matches across species. Chromosome heights are normalized to equal heights. See Fig. 3 for further layout interpretation.

| P. waltl (query) | P. waltl (subject) | | | | | | |
|---|---|---|---|---|---|---|---|
| Gene | Gene | Start | End | Chrm # | Coverage | # HSPs | Chrm |
| AIRE_nanopore | Gene_3 | 504063889 | 504201253 | 11 | 2570/2570 (100.0%) | 88 | |
| | Gene_15 | 979825436 | 979825708 | 6 | 271/2570 (10.5%) | 4 | |
| | Gene_2 | 476873610 | 477290777 | 11 | 258/2570 (10.0%) | 2 | |
| | Gene_6 | 804609142 | 804609394 | 11 | 253/2570 (9.8%) | 4 | |
| | Gene_59 | 1062468268 | 1062468510 | 10 | 246/2570 (9.6%) | 4 | |
| | Gene_72 | 2316932354 | 2316932588 | 1_2 | 244/2570 (9.5%) | 5 | |
| | Gene_70 | 1716638913 | 1716639164 | 1_2 | 238/2570 (9.3%) | 3 | |
| | Gene_75 | 1095782727 | 1095782988 | 8 | 231/2570 (9.0%) | 4 | |
| | Gene_62 | 391569872 | 391570116 | 5 | 228/2570 (8.9%) | 8 | |
| | Gene_24 | 681706665 | 681785069 | 12 | 225/2570 (8.8%) | 3 | |
| | Gene_61 | 180093397 | 180093615 | 5 | 223/2570 (8.7%) | 4 | |
| | Gene_43 | 45856306 | 45856527 | 7 | 222/2570 (8.6%) | 1 | |
| | Gene_51 | 1373041773 | 1373041985 | 4_2 | 213/2570 (8.3%) | 2 | |
| | Gene_76 | 37893042 | 37893288 | 9 | 213/2570 (8.3%) | 5 | |
| | Gene_9 | 171053143 | 171053370 | 6 | 211/2570 (8.2%) | 3 | |
| | Gene_69 | 1285765167 | 1285765422 | 1_2 | 207/2570 (8.1%) | 3 | |
| | Gene_57 | 1028054240 | 1028054441 | 10 | 205/2570 (8.0%) | 3 | |
| | Gene_35 | 798636272 | 798636385 | 3_1 | 204/2570 (7.9%) | 2 | |
| | Gene_77 | 422334996 | 422335294 | 9 | 202/2570 (7.9%) | 3 | |
| | Gene_74 | 419778857 | 419779079 | 8 | 201/2570 (7.8%) | 4 | |
| | Gene_63 | 1609440069 | 1609440259 | 5 | 198/2570 (7.7%) | 2 | |
| | Gene_33 | 400355946 | 400356161 | 3_1 | 196/2570 (7.6%) | 4 | |
| | Gene_60 | 984158088 | 984158307 | 1_1 | 196/2570 (7.6%) | 3 | |
| | Gene_71 | 2195043835 | 2195044050 | 1_2 | 195/2570 (7.6%) | 3 | |
| | Gene_73 | 51239676 | 51239912 | 8 | 188/2570 (7.3%) | 3 | |
| | Gene_12 | 825551622 | 825551804 | 6 | 183/2570 (7.1%) | 1 | |
| | Gene_64 | 1840150846 | 1840151053 | 2_2 | 183/2570 (7.1%) | 3 | |
| | Gene_56 | 967281061 | 967281234 | 10 | 177/2570 (6.9%) | 2 | |
| | Gene_17 | 1051500974 | 1051517126 | 6 | 171/2570 (6.7%) | 2 | |
| | Gene_32 | 244101494 | 244101657 | 3_1 | 171/2570 (6.7%) | 2 | |
| | Gene_10 | 586487743 | 586487907 | 6 | 168/2570 (6.5%) | 2 | |
| | Gene_1 | 43349034 | 43349198 | 11 | 165/2570 (6.4%) | 1 | |
| | Gene_27 | 227387330 | 227387494 | 4_1 | 165/2570 (6.4%) | 2 | |
| | Gene_46 | 912709490 | 912709646 | 7 | 165/2570 (6.4%) | 2 | |
| | Gene_68 | 2196067479 | 2196067638 | 3_2 | 163/2570 (6.3%) | 2 | |
| | Gene_45 | 537206892 | 537207052 | 7 | 162/2570 (6.3%) | 2 | |
| | Gene_78 | 1148739683 | 1148739841 | 9 | 162/2570 (6.3%) | 2 | |
| | Gene_19 | 1507077937 | 1507078095 | 6 | 156/2570 (6.1%) | 2 | |
| | Gene_48 | 1120803723 | 1120803878 | 7 | 156/2570 (6.1%) | 1 | |
| | Gene_44 | 87705901 | 87706050 | 7 | 150/2570 (5.8%) | 1 | |
| | Gene_47 | 1048451174 | 1048451323 | 7 | 150/2570 (5.8%) | 1 | |
| | Gene_52 | 2005600 | 2005743 | 10 | 144/2570 (5.6%) | 1 | |
| | Gene_31 | 23071676 | 23071813 | 3_1 | 138/2570 (5.4%) | 1 | |
| | Gene_11 | 659250999 | 659251136 | 6 | 138/2570 (5.4%) | 1 | |
| | Gene_28 | 260294731 | 260294868 | 4_1 | 138/2570 (5.4%) | 1 | |
| | Gene_36 | 1094214776 | 1094214907 | 3_1 | 132/2570 (5.1%) | 1 | |
| | Gene_16 | 1015984541 | 1015984666 | 6 | 126/2570 (4.9%) | 1 | |
| | Gene_67 | 2079614177 | 2079614302 | 3_2 | 126/2570 (4.9%) | 1 | |
| | Gene_20 | 155653683 | 155653799 | 12 | 117/2570 (4.6%) | 1 | |
| | Gene_50 | 1054080249 | 1054080365 | 4_2 | 117/2570 (4.6%) | 1 | |
| | Gene_55 | 629103547 | 629103663 | 10 | 117/2570 (4.6%) | 1 | |
| | Gene_25 | 179236083 | 179236196 | 4_1 | 114/2570 (4.4%) | 1 | |
| | Gene_26 | 201227981 | 201228094 | 4_1 | 114/2570 (4.4%) | 1 | |
| | Gene_18 | 1096172014 | 1096172121 | 6 | 108/2570 (4.2%) | 1 | |
| | Gene_54 | 139553202 | 139553306 | 10 | 105/2570 (4.1%) | 1 | |
| | Gene_4 | 546393916 | 546394017 | 11 | 102/2570 (4.0%) | 1 | |
| | Gene_37 | 1230728824 | 1230728925 | 3_1 | 102/2570 (4.0%) | 1 | |
| | Gene_29 | 957640618 | 957640719 | 4_1 | 102/2570 (4.0%) | 1 | |
| | Gene_7 | 931487736 | 931487834 | 11 | 99/2570 (3.9%) | 1 | |
| | Gene_38 | 1629699778 | 1629699873 | 3_1 | 96/2570 (3.7%) | 1 | |
| | Gene_66 | 2060186370 | 2060186465 | 3_2 | 96/2570 (3.7%) | 1 | |
| | Gene_23 | 659377086 | 659377178 | 12 | 93/2570 (3.6%) | 1 | |
| | Gene_39 | 1823627396 | 1823627488 | 3_1 | 93/2570 (3.6%) | 1 | |
| | Gene_42 | 1942894742 | 1942894834 | 3_1 | 93/2570 (3.6%) | 1 | |
| | Gene_5 | 626741243 | 626741329 | 11 | 87/2570 (3.4%) | 1 | |
| | Gene_34 | 620107975 | 620108061 | 3_1 | 87/2570 (3.4%) | 1 | |
| | Gene_58 | 1034640554 | 1034640640 | 10 | 87/2570 (3.4%) | 1 | |
| | Gene_41 | 1938903242 | 1938903325 | 3_1 | 84/2570 (3.3%) | 1 | |
| | Gene_49 | 1130893399 | 1130893482 | 7 | 84/2570 (3.3%) | 1 | |
| | Gene_8 | 970265395 | 970265478 | 11 | 84/2570 (3.3%) | 1 | |
| | Gene_22 | 366192317 | 366192397 | 12 | 81/2570 (3.2%) | 1 | |
| | Gene_40 | 1903604053 | 1903604133 | 3_1 | 81/2570 (3.2%) | 1 | |
| | Gene_13 | 870183533 | 870183613 | 6 | 81/2570 (3.2%) | 1 | |
| | Gene_14 | 876212080 | 876212160 | 6 | 81/2570 (3.2%) | 1 | |
| | Gene_65 | 2019497488 | 2019497568 | 3_2 | 81/2570 (3.2%) | 1 | |
| | Gene_30 | 1004309743 | 1004309823 | 4_1 | 81/2570 (3.2%) | 1 | |
| | Gene_53 | 71129240 | 71129317 | 10 | 78/2570 (3.0%) | 1 | |
| | Gene_21 | 222501193 | 222501267 | 12 | 75/2570 (2.9%) | 1 | |

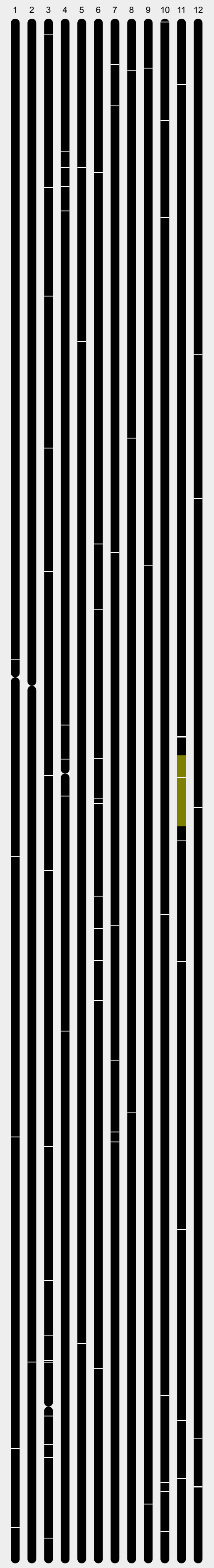

**Supplemental Figure 4. Complete tBLASTx search results for AIRE gene signal detection in *P. waltl*.** See Figure 9C description for details.

| H. sapiens | | | | | P. waltl | | | | | | |
|---|---|---|---|---|---|---|---|---|---|---|---|
| **#** | **Gene** | **Start** | **End** | **Chrm** | **Gene** | **Start** | **End** | **Chrm #** | **Coverage** | **# HSPs** | **Chrm** |
| 1 | RBL1 | 36996349 | 37095997 | 20 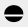 | Gene_7 | 306130219 | 306446903 | 12 | 1722/2420 (71.2%) | 18 | 8 12 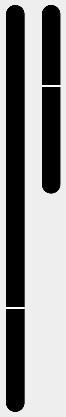 |
| | | | | | Gene_4 | 306006516 | 306446903 | 12 | 2015/3233 (62.3%) | 21 | |
| | | | | | Gene_6 | 305922326 | 306430662 | 12 | 2117/5469 (38.7%) | 21 | |
| | | | | | Gene_10 | 305922326 | 306430662 | 12 | 2117/5469 (38.7%) | 21 | |
| | | | | | Gene_3 | 305922326 | 306446903 | 12 | 2183/5686 (38.4%) | 22 | |
| | | | | | Gene_11 | 305922326 | 306430662 | 12 | 2052/5404 (38.0%) | 20 | |
| | | | | | Gene_5 | 305922326 | 306446903 | 12 | 2078/5521 (37.6%) | 21 | |
| | | | | | Gene_9 | 305922326 | 306446903 | 12 | 2078/5521 (37.6%) | 21 | |
| | | | | | Gene_2 | 305922326 | 306446903 | 12 | 2183/5824 (37.5%) | 22 | |
| | | | | | Gene_1 | 305922326 | 306446903 | 12 | 2118/5759 (36.8%) | 21 | |
| | | | | | Gene_8 | 1147971785 | 1147971937 | 8 | 153/2420 (6.3%) | 1 | |

**Supplemental Figure 5. Genomic screening for *Rbl1* signals in *P. waltl* using human *RBL1* as reference sequence.** tBLASTx search results against the *P. waltl* genome visualized with *Novabrowse*. “Start” and “End” columns indicate chromosomal positions. Chromosome sizes are scaled proportionally to their actual lengths. "# HSPs" column shows the number of High-scoring Segment Pairs joined to form each putative gene signal. See Fig. 3 for further layout interpretation.

| # | Tool | Feature | Verdict | Description |
|---|---|---|---|---|
| 1 | NCBI BLAST | Coordinate-based genomic region search | NO | Accepts sequences/accessions only; cannot input chromosomal coordinates to extract genes from a region for BLAST querying. https://blast.ncbi.nlm.nih.gov/doc/blast-topics/queryinpanddatasel.html |
| 2 | NCBI BLAST | Distance-based HSP clustering | NO | HSPs reported individually; no user-configurable distance-based clustering into putative gene units. Johnson et al. 2008, NAR. https://pmc.ncbi.nlm.nih.gov/articles/PMC2447716/ |
| 3 | NCBI BLAST | Multi-species gene synteny visualization | NO | No synteny visualization of any kind. https://blast.ncbi.nlm.nih.gov/doc/blast-topics/resultformatoptions.html |
| 4 | NCBI BLAST | Chromosomal visualization & mapping | NO | BLAST links to NCBI Genome Data Viewer (GDV) which shows hit locations as arrowheads on chromosome ideograms, but this is a separate tool, not integrated into BLAST output itself. https://ncbiinsights.ncbi.nlm.nih.gov/2017/10/24/ncbis-genome-data-viewer-gdv-to-replace-map-viewer/ |
| 5 | NCBI BLAST | Coverage visualization | YES | Graphic Summary shows HSP positions on query axis colored by alignment score (bit score), less informative compared to Novabrowse. https://blast.ncbi.nlm.nih.gov/doc/blast-quick-start-results/graphicsummarytab.html |
| 6 | NCBI BLAST | Integrated BLAST search | YES | Core function: runs BLASTN, BLASTP, BLASTX, TBLASTN, TBLASTX. Johnson et al. 2008, NAR. https://pmc.ncbi.nlm.nih.gov/articles/PMC2447716/ |
| 7 | NCBI BLAST | Custom BLAST database support | NO | Limited to NCBI's pre-built databases (nr, nt, RefSeq, etc.). There's a "blast2seq" feature for pasting a small subject sequence, but it can't handle genome-scale assemblies. |
| 8 | NCBI BLAST | Isoform-aware hit consolidation | NO | Each isoform hit reported as separate alignment; no gene-level grouping with expandable transcript detail. https://blast.ncbi.nlm.nih.gov/doc/blast-topics/resultformatoptions.html |
| 1 | Ensembl BLAST | Coordinate-based genomic region search | NO | BLAST/BLAT input accepts sequence only; coordinate-based retrieval requires separate REST API. http://www.ensembl.org/Help/View?id=451 |
| 2 | Ensembl BLAST | Distance-based HSP clustering | NO | Individual HSPs reported separately; no distance-based clustering into gene units. http://www.ensembl.org/Help/View?id=451 |
| 3 | Ensembl BLAST | Multi-species gene synteny visualization | NO | Synteny view exists as a separate tool, is strictly pairwise (two genomes only) and is functionally disconnected from BLAST results data: https://www.ensembl.org/info/genome/compara/synteny.html |
| 4 | Ensembl BLAST | Chromosomal visualization & mapping | YES | BLAST results mapped onto karyotype ideogram with colored arrowheads showing hit locations across chromosomes. http://www.ensembl.org/Help/View?id=451 |
| 5 | Ensembl BLAST | Coverage visualization | YES | Not directly visible from BLAST results view (is 1 page away and visible only per hit). Query sequence HSP distribution shown as colored boxes (black/white/red), but not identity-percent-coded coverage bars (green/yellow/orange/red gradient). http://www.ensembl.org/Help/View?id=451 |
| 6 | Ensembl BLAST | Integrated BLAST search | YES | Runs BLASTN, BLASTX, TBLASTX, and BLAT natively. Martin et al. 2023, NAR. https://pmc.ncbi.nlm.nih.gov/articles/PMC10767893/ |
| 7 | Ensembl BLAST | Custom BLAST database support | NO | Strictly limited to Ensembl's curated species databases. |
| 8 | Ensembl BLAST | Isoform-aware hit consolidation | NO | Each transcript isoform appears as a separate hit entry; no gene-level consolidation with expandable isoform detail. http://www.ensembl.org/Help/View?id=451 |
| 1 | UCSC (BLAT) | Coordinate-based genomic region search | NO | Browser navigates to coordinates, but does not extract genes from a genomic region and use them as BLAST queries. Coordinate navigation is genome browsing, not coordinate-based search for gene discovery. https://genome.ucsc.edu/goldenPath/help/hgTracksHelp.html |
| 2 | UCSC (BLAT) | Distance-based HSP clustering | NO | BLAT internally merges HSPs with a hardcoded ~300bp gap threshold; this is not user-configurable distance-based clustering for gene unit discovery. Kent 2002, Genome Res. https://genome.cshlp.org/content/12/4/656 |
| 3 | UCSC (BLAT) | Multi-species gene synteny visualization | NO | Chain/Net and Snake tracks are strictly pairwise (two genomes); no Multi-species gene synteny visualization with interactive ribbons connecting 3+ species. https://genome.ucsc.edu/goldenPath/help/hgTracksHelp.html |
| 4 | UCSC (BLAT) | Chromosomal visualization & mapping | YES | Chromosome ideogram shows position for the current viewed region only; BLAT results list does not provide a chromosomal overview mapping all hits simultaneously. https://pmc.ncbi.nlm.nih.gov/articles/PMC4101998/ |
| 5 | UCSC (BLAT) | Coverage visualization | NO | No query-centric identity-percent-coded coverage bars. BLAT shows tabular match/mismatch statistics and base-level coloring in alignment detail, but no graphical coverage visualization. https://genome.ucsc.edu/FAQ/FAQblat.html |
| 6 | UCSC (BLAT) | Integrated BLAST search | YES | Though it uses BLAT (not BLAST); . Kent 2002. https://genome.cshlp.org/content/12/4/656 |
| 7 | UCSC (BLAT) | Custom BLAST database support | NO | Limited to UCSC's pre-built assemblies. Technically possible via Assembly Hubs, but that requires running your own gfServer infrastructure |
| 8 | UCSC (BLAT) | Isoform-aware hit consolidation | NO | BLAT returns independent per-query alignments with no gene-level grouping. https://genome.ucsc.edu/goldenpath/help/blatSpec.html |
| 1 | SequenceServer | Coordinate-based genomic region search | NO | Accepts only raw sequences pasted or uploaded; no coordinate-based gene extraction. Priyam et al. 2019, MBE. https://pmc.ncbi.nlm.nih.gov/articles/PMC6878946/ |
| 2 | SequenceServer | Distance-based HSP clustering | NO | Individual HSPs displayed as separate entries; no distance-based clustering. https://sequenceserver.com/blog/visualizing_blast_results/ |
| 3 | SequenceServer | Multi-species gene synteny visualization | NO | No synteny visualization; enterprise version has custom add-ons but not standard synteny. https://sequenceserver.com/cloud/ |
| 4 | SequenceServer | Chromosomal visualization & mapping | NO | No chromosome ideograms or chromosomal mapping; links to external genome browsers. https://sequenceserver.com/blast-interface/ |
| 5 | SequenceServer | Coverage visualization | YES | Alignment overview shows HSP positions on query colored by E-value (not identity-percent gradient). Circos-style length distribution plots available. https://sequenceserver.com/blog/visualizing_blast_results/ |
| 6 | SequenceServer | Integrated BLAST search | YES | Core function: provides web GUI for NCBI BLAST+. Priyam et al. 2019, MBE. https://pmc.ncbi.nlm.nih.gov/articles/PMC6878946/ |
| 7 | SequenceServer | Custom BLAST database support | YES | Supports user-provided genome assemblies as search targets. |
| 8 | SequenceServer | Isoform-aware hit consolidation | NO | Each hit displayed individually; no gene-level grouping. https://pmc.ncbi.nlm.nih.gov/articles/PMC6878946/ |
| 1 | MCScanX | Coordinate-based genomic region search | NO | Requires whole-genome GFF + BLASTP input files; no coordinate-based region search. Wang et al. 2012, NAR. https://pmc.ncbi.nlm.nih.gov/articles/PMC3326336/ |
| 2 | MCScanX | Distance-based HSP clustering | NO | Operates at gene level (collinear gene pairs), not HSP level; . Wang et al. 2012. https://pmc.ncbi.nlm.nih.gov/articles/PMC3326336/ |
| 3 | MCScanX | Multi-species gene synteny visualization | YES | circle_plotter supports 3+ species on a single plot, dual_synteny is pairwise only. Wang et al. 2012, Fig 3. https://pmc.ncbi.nlm.nih.gov/articles/PMC3326336/ |
| 4 | MCScanX | Chromosomal visualization & mapping | YES | Four visualization programs (dot_plotter, circle_plotter, dual_synteny, bar_plotter) map syntenic blocks onto chromosome representations. Wang et al. 2012, Fig 3. https://pmc.ncbi.nlm.nih.gov/articles/PMC3326336/ |
| 5 | MCScanX | Coverage visualization | NO | No query-centric coverage visualization; provides syntenic block depth counts only. Wang et al. 2012, Fig 2. https://pmc.ncbi.nlm.nih.gov/articles/PMC3326336/ |
| 6 | MCScanX | Integrated BLAST search | NO | Requires pre-computed external all-vs-all BLASTP results as input; does not run BLAST itself. Wang et al. 2012. https://pmc.ncbi.nlm.nih.gov/articles/PMC3326336/ |
| 7 | MCScanX | Custom BLAST database support | YES | Supports user-provided genome assemblies as search targets. |
| 8 | MCScanX | Isoform-aware hit consolidation | NO | Expects single representative protein per gene in input; no isoform handling. Wang et al. 2012. https://pmc.ncbi.nlm.nih.gov/articles/PMC3326336/ |

**Supplemental Table 1. Detailed evidence and references supporting the feature comparison verdicts in Table 1.** Each entry documents the specific basis for the YES or NO verdict assigned to a given tool-feature combination, including relevant documentation.

| Protein | Protein name | Isoform |
|---|---|---|
| XP_069069929.1 | nuclear body protein SP140-like protein | X1 |
| XP_069069930.1 | nuclear body protein SP140-like protein | X1 |
| XP_069069931.1 | nuclear body protein SP140-like protein | X2 |
| XP_069069932.1 | nuclear body protein SP140-like protein | X3 |
| XP_069069933.1 | nuclear body protein SP140-like protein | X3 |
| XP_069069934.1 | sp110 nuclear body protein-like | X4 |
| XP_069069935.1 | sp110 nuclear body protein-like | X5 |

**Supplemental Table 2. Protein isoforms of *P. waltl Loc138265795*.** NCBI-annotated protein accessions, names, and isoform designations for all seven protein variants encoded by *Loc138265795*, identified as the highest-scoring tBLASTx match to both human *AIRE* and *X. tropicalis aire*.
